# Modifications in the T arm of tRNA globally determine tRNA maturation, function and cellular fitness

**DOI:** 10.1101/2023.11.11.566336

**Authors:** Sarah K. Schultz, Christopher D. Katanski, Mateusz Halucha, Noah Pena, Richard P. Fahlman, Tao Pan, Ute Kothe

## Abstract

All elongator tRNAs harbor 5-methyluridine 54 and pseudouridine 55 in the T arm, which are generated by the enzymes TrmA and TruB, respectively. *Escherichia coli* TrmA and TruB both act as tRNA chaperones, and strains lacking *trmA* or *truB* are outcompeted by wildtype. Here, we investigate how TrmA and TruB contribute to cellular fitness. Deletion of *trmA* and *truB* in *E. coli* causes a global decrease in aminoacylation and alters other tRNA modification such as acp^3^U47. Whereas overall protein synthesis is not affected in Δ*trmA* and Δ*truB* strains, the translation of a specific subset of codons is significantly impaired, and the expression of many specific proteins is translationally changed. In conclusion, we demonstrate that universal modifications of the tRNA T arm are critical for global tRNA function by enhancing tRNA maturation, tRNA aminoacylation, and translation, thereby improving cellular fitness and explaining *trmA* and *truB* conservation.

## Introduction

Transfer RNAs (tRNAs) are the most densely and most diversely modified class of RNA, and all organisms invest significant energy into synthesizing large numbers of tRNA modifying enzymes ^1^. In humans, defects in tRNA modification enzymes are frequently associated with diseases including cancer, neurological disorders, and mitochondrial diseases ^2^. Within tRNAs, two general clusters of modifications are apparent. Modifications in the anticodon stem-loop generally play direct and essential roles during translation; however, functional roles for modifications within the tRNA elbow oftentimes remain elusive ^3,4^. Two of the most common and conserved tRNA modifications are 5-methyluridine (m^5^U) 54 and pseudouridine (Ψ) 55 (Figure 1). These modifications are found at adjacent positions within the T arm of nearly every elongator tRNA throughout all domains of life. In *Escherichia coli*, m^5^U54 is formed by the methyltransferase TrmA ^5^, whereas Ψ55 is generated by the pseudouridine synthase TruB ^6,7^. Like their modifications, the TrmA and TruB enzymes are highly conserved throughout all domains of life, although some archaea and diverse bacteria utilize alternative enzymatic strategies to introduce m^5^U54 and Ψ55 ^4^.

**Figure 1.**
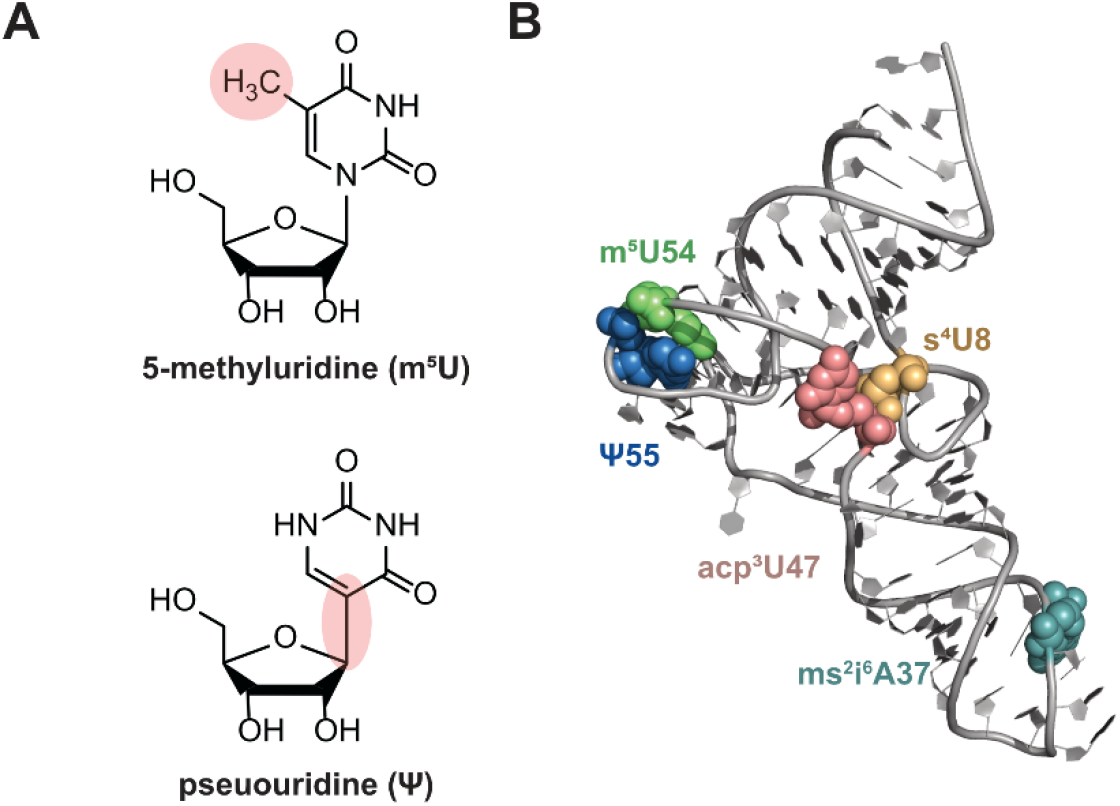
Structures and location of m^5^U54 and Ψ55. **(A)**Chemical structures of m^5^U (top) and Ψ (bottom) nucleosides. The region of the modification is highlighted in red. **(B)** Location of m^5^U54 (green spheres), Ψ55 (blue spheres), s^4^U8 (yellow spheres), acp^3^U47 (pink spheres), and ms^2^i^6^A37 (teal spheres) within the crystal structure of unmodified *E. coli* tRNA^Phe^ (PDB 3L0U) ^48^.

Highlighting the importance of TrmA and TruB and their respective modifications, m^5^U54 and Ψ55 are the only two modifications found within every *E. coli* tRNA ^8^. In this model organism, the *trmA* and *truB* genes are non-essential under ideal conditions but are important for bacterial fitness in co-culture experiments ^9-12^. In addition to modifying tRNAs, TrmA and TruB act as tRNA chaperones, folding tRNA independently of their modification activity ^10,11,13^. The tRNA chaperone activity of TruB is responsible for optimizing fitness in *E. coli* ^9,11^, whereas TruB pseudouridylation is important for growth at cold temperatures in *Thermus thermophilus* ^14^,heat shock recovery in *E. coli* ^15^, and *Shigella flexneri* virulence ^16^. Both the chaperone activity and modification activities of TrmA play roles in *E. coli* fitness ^10^, but the methylation activity seems dispensable in yeast ^12,17^. Finally, the m^5^U54 and Ψ55 modifications themselves stabilize tRNA ^3^. Taken together, the conserved enzymes TrmA and TruB are bifunctional enzymes, playing two distinct roles during tRNA maturation by folding tRNA and introducing tRNA-stabilizing modifications. Supporting the importance for TrmA and TruB homologs in humans, the TrmA homolog TRMT2A is a potential biomarker for the recurrence of certain cancers and is a potential target for polyQ diseases ^18,19^. Moreover, the TruB homologs, TRUB1/2, are linked to maternally inherited diabetes through a mitochondrial tRNA^Glu^ mutation that prevents tRNA pseudouridylation ^20,21^. Contrasting the conservation of TrmA and TruB enzymes, their implications in human diseases, and abundance of m^5^U54 and Ψ55 modifications, only a handful of subtle growth phenotypes have been identified for model organisms lacking these enzymes ^15,16^.

Here we address the paradox of almost all cells investing energy into modifying and folding the majority of tRNAs by TrmA and TruB, although these enzymes seem to be unimportant for growth during ideal and various stress conditions. We examined which functional tRNA processes are affected in the absence of *trmA* and *truB* genes by measuring tRNA abundance, tRNA modification, tRNA aminoacylation, as well as codon-specific and global protein translation within *E. coli ΔtrmA* and *ΔtruB* knockout strains. Thereby, this work highlights that absence of TrmA or TruB results in several defects during tRNA maturation, aminoacylation and protein translation, explaining why cells lacking TrmA or TruB exhibit reduced fitness.

## Results

### tRNA T arm modification globally enhances aminoacylation

To dissect the roles of TruB and TrmA for tRNA function and protein synthesis, we used *E. coli ΔtruB* and *ΔtrmA* strains from the Keio Collection ^22^ in addition to preparing a double-knockout strain lacking both *truB* and *trmA* genes. Previously, TrmA and TruB chaperone activity has been assessed *in vitro* by detecting tRNA aminoacylation (charging) because only correctly folded tRNA is efficiently charged ^10,11,23^. Therefore, we hypothesized that the deletion of *trmA* and/or *truB* increases the fraction of misfolded tRNA and in turn decreases the aminoacylation levels of some or all tRNAs *in vivo*. To quantitatively compare both the abundance and the aminoacylation levels for all tRNAs, we used multiplex small RNA sequencing (MSR-seq) which simultaneously measures tRNA abundance, modification, and charging ^24^. Herein, the 3′ ribose of uncharged tRNA is oxidized leading to β-elimination of the 3′ adenine of the tRNA whereas the terminal 3′ adenine is protected in aminoacylated tRNA. After Illumina sequencing, comparison of sequences ending in 3′ CC (deacylated) to 3′ CCA (aminoacylated) allows for quantification of the aminoacylation level for individual tRNA isoacceptors while also providing information about relative tRNA abundance and modification content ^24,25^.

First, we asked whether TrmA and/or TruB and their resulting modifications m^5^U54 and Ψ55 promote the cellular stability and thereby steady-state abundance of tRNAs *in vivo* as observed for other tRNA modifying enzymes ^1,26,27^. Within the single Δ*trmA* and Δ*truB* strains, no tRNA is decreased or increased more than 1.5-fold upon modification enzyme knockout (Fig. 2A, Fig. 2B, Extended Data Fig. 1). In the double-knockout strain, only two tRNAs, tRNA^Leu^TAG and tRNA^Leu^CAG-2-1 are downregulated by ∼1.5-fold compared to the wildtype strain (Fig. 2C). The decreased abundances for these two tRNAs may be associated with lower stability of their stem-loop in the variable region compared to closely related tRNA^Leu^GAG and tRNA^Leu^CAG-1-1 (Fig. 2D). These results indicate that the lack of TrmA and TruB has only minimal effects on the overall cellular stability of tRNAs in *E. coli*.

**Fig. 2.**
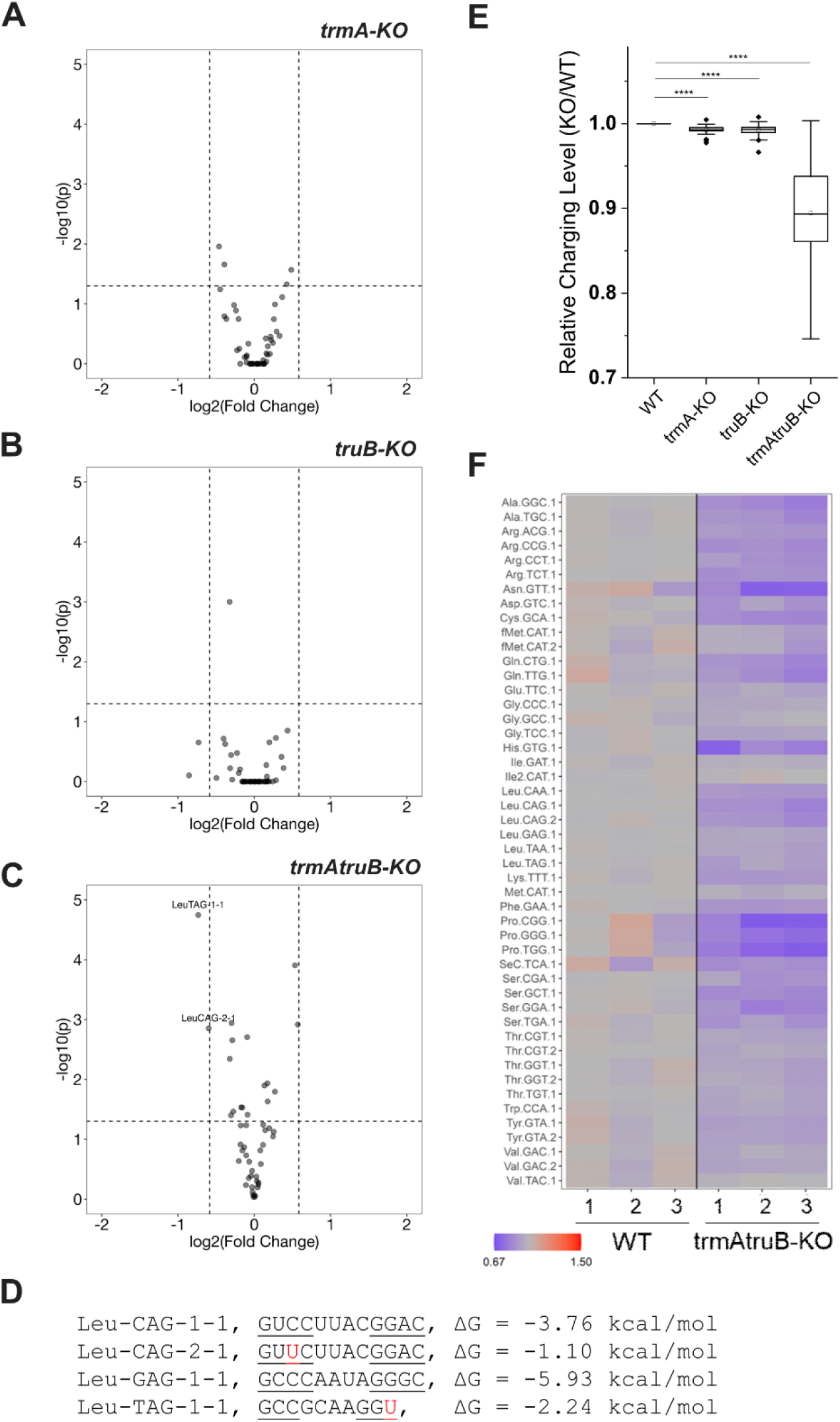
tRNA aminoacylation is reduced globally in *trmA* and *truB* knockout strains. tRNA abundance change in single and double knockout strains depicted as volcano plots. Vertical dashed lines indicate threshold change of 1.5-fold, horizontal dashed lines indicate *p* = 0.05. **(A)** *ΔtrmA*; **(B)** *ΔtruB*; **(C)** *ΔtrmAΔtruB*. **(D)** Variable region stem-loop sequences and free energy calculations of tRNA^Leu^(xAG). U of G-U pair is shown in red. **(E)** Charging level change of all tRNAs in wild-type, *ΔtrmA*; *ΔtruB*; and *ΔtrmAΔtruB* strains. Box and whisker plot, **** *p* < 10^-4^. **(F)** Heatmap of charging level change for each tRNA in *ΔtrmAΔtruB* over *E. coli* BW25113 wildtype (WT) showing each biological replicate.

Next, we examined changes in aminoacylation upon *trmA* or *truB* knockout. Comparison of aminoacylation levels revealed a significant global decrease in tRNA charging in the Δ*truB* and Δ*trmA* strains, with more pronounced effects in the double Δ*trmA*Δ*truB* strain (Fig. 2E-F). In the double knockout strain, tRNA aminoacylation is on average 10 % lower than in the wildtype strain, suggesting additive effects of removing both TrmA and TruB (Fig. 2E). Almost all individual tRNAs are less charged, but changes in charging level are variable (Fig. 2F). This data supports our hypothesis that TrmA and TruB affect tRNA charging of almost all tRNAs *in vivo*, and the effect is much more pronounced upon losing both modifications.

### m^5^U54 and Ψ55 influence the modification of several other tRNA nucleotides

Since certain tRNA modifications, such as 3-(3-amino-3-carboxypropyl)uridine at position 47 in the tRNA variable loop (acp^3^U47, Fig. 3A), cause mutations or deletions during reverse transcription when a thermostable reverse transcriptase (Superscript IV) is used for MSR-seq, the tRNA sequencing dataset also enables detecting changes in mutation and/or deletion frequencies at specific nucleotides that reflect tRNA modifications ^24^. Comparing the *ΔtrmA* strain to wild-type *E. coli*, we observe reduced mutation and deletion frequencies at position 47 within the majority of tRNAs that carry the acp^3^U47 modification, suggesting that the acp^3^U47 modification is less abundant in the absence of TrmA/m^5^U54 (Fig. 3B, Fig. 3C). Similar, further reduced mutation and deletion frequencies at U47 are observed for the *ΔtrmAΔtruB* strain; however, single knockout of *truB* does not affect acp^3^U47 content in most tRNAs with the exception of tRNA^Phe^ (Fig 3C). To validate the acp^3^U47 content in the absence of TrmA and TruB, we conducted primer extension analysis using a reverse transcriptase with low processivity (AMV RT) for tRNA extracted from *E. coli* observing the reaction stop at the bulky modification acp^3^U47 (Fig. 3D); in the absence of this modification, the primer is extended either to the 5′ end of the tRNA or to the next bulky tRNA modification (ms^2^i^6^A37 in the case of tRNA^Phe^GAA). Determining the ratio of band intensities for primer extension stops at position 47 to 37 confirms a reduction of acp^3^U47 in tRNA^Phe^GAA upon deletion of *trmA, truB*, or both (Fig. 3D, Extended Data Fig. 2A-C). Again, TrmA seems to play a larger role in acp^3^U47 formation than TruB (Fig. 3C, Fig. 3D).

**Fig. 3.**
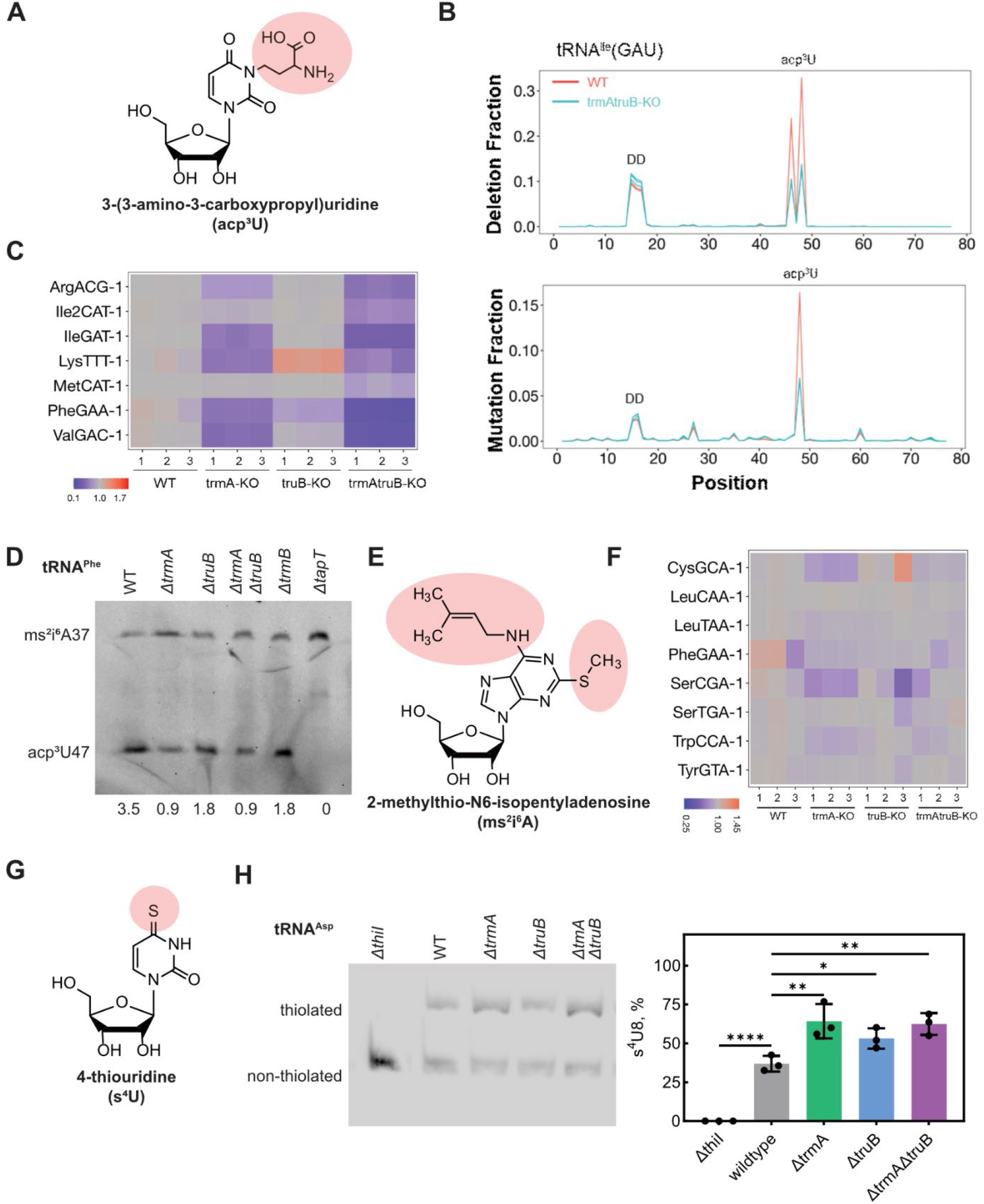
Deletion of *trmA* and/or *truB* changes abundance of additional tRNA modifications. **(A)**Chemical structure of acp^3^U with modified portions highlighted in red. **(B)** Mutation and deletion fraction of tRNA^Ile^(GAU) reads in WT and *ΔtrmAΔtruB* strains. Known modifications for this tRNA include acp^3^U47 which generates strong mutation and deletion signatures, and consecutive dihydrouridines (D) D19D20 which also generate mutation and deletion signatures. *ΔtrmAΔtruB* reduces the mutation and deletion fraction of acp^3^U47, but not those of D19D20. **(C)** Heatmap of acp^3^U47 level change of *ΔtrmA* and/or *ΔtruB* over WT showing each biological replicate. All 7 *E. coli* tRNAs with known acp^3^U47 modifications are shown. Data shown are the sum of mutation+deletion fraction at corresponding U47 position relative to WT. Blue indicates a decrease of acp^3^U in the double-KO strain. **(D)** Validation of a decrease in acp^3^U47 content upon deletion of either *trmA* or *truB* by tRNA^Phe^ primer extension and reverse transcription (RT). Ratios of acp^3^U47/ms^2^i
^6^A37 intensities are indicated at the bottom of the gel. Gel displayed is representative of two biological replicates. All uncropped gels and quantifications are displayed in Extended Data Figure 2. **(E)** Chemical structure of ms^2^i^6^A37 with modified portions highlighted in red. **(F)** Heatmap of mutation and deletion frequency changes at position 37 for all *E. coli* tRNAs harboring a ms^2^i^6^A37 modification. **(G)** Chemical structures of s^4^U with modified portions highlighted in red. **(H)** The presence of s^4^U8 in knockout strains grown in LB medium containing 2% formate was detected by APM-PAGE followed by Northern Blotting for tRNA^Asp^ (left) and quantified based on the band intensities of the thiolated and non-thioated form (right). **** indicates *p* < 10^-4^, ** indicates *p* < 0.01, and * indicates *p* <0.05. Three biological replicates were performed, and all replicates are shown in Extended Data Figure 3.

In addition to this change in the tRNA variable loop, the tRNA sequencing data also indicate potential changes in the anticodon arm. Among these, we noticed several changes at the conserved purine 37 adjacent to the anticodon, which is modified to ms^2^i^6^A in several tRNAs (Fig. 3E). Particularly in the absence of *trmA*, the deletion/mutation signature at A37 is changed relative to wildtype within several tRNAs known to contain ms^2^i^6^A (tRNA^Leu^TAA, tRNA^Phe^GAA, tRNA^Trp^CCA, tRNA^Cys^GCA, and tRNA^Ser^CGA); however, other ms^2^i^6^A37-containing tRNAs may not be affected (tRNA^Tyr^GTA, tRNA^Leu^CAA, Fig. 3F). A smaller effect on ms^2^i^6^A37 is observed in the *ΔtruB* and double knockout strains. Only *E. coli* tRNA^Arg^ACG contains the edited base inosine 34, which generates a mutation during reverse transcription. The absence of *trmA* and/or *truB* does not affect tRNA deamination, as all strains had a high level of mutation (>97%) at A34 of tRNA^Arg^ACG (Extended Data Fig. 2D).

Finally, we also wondered if other common modifications closer to m^5^U54 and Ψ55 within the tRNA tertiary structure are affected by the loss of TrmA and/or TruB. In particular, we examined whether s^4^U8 (Fig. 3G) abundance is dependent on presence of TrmA or TruB by using urea-PAGE containing a phenylmercury compound that decreases the migration velocity of sulfur-containing tRNAs, wherein we detected specific tRNAs by Northern blotting ^28^. For many tRNAs, a high level (near 100%) of tRNA thiolation is observed in the absence and presence of TrmA and TruB enzymes (Extended Data Fig. 3). In contrast, tRNA^Asp^ was only ∼30-40 % thiolated in the wildtype strain. Interestingly, the proportion of thiolated tRNA^Asp^ increased to ∼50 % in the *ΔtruB* strain and to ∼60 % in the *ΔtrmA* and *ΔtrmAΔtruB* strains (Fig. 3H), suggesting the absence of TrmA and TruB promotes s^4^U8 formation by ThiI. In summary, TruB and TrmA affect other tRNA modifications at several positions including U8, A37 and U47 (and possibly more that do not provide strong reverse transcription signatures in sequencing) thereby influencing several steps during tRNA maturation.

### TruB and TrmA affect translation of specific codons

Since tRNA aminoacylation and modification are affected in the absence of TruB and TrmA, we next investigated how cellular protein synthesis is impacted. To assess potential effects on global translation, we utilized biorthogonal noncanonical amino acid tagging (BONCAT), wherein L-azidohomoalanine (L-AHA) is incorporated into newly synthesized proteins instead of L-methionine allowing for labelling with a rhodamine dye^29^. Overall quantification of new proteins does not reveal statistically significant changes between *E. coli* wild-type and the Δ*trmA/*Δ*truB* strains (Fig. 4A, Extended Data Fig. 4). However, upon careful inspection of the SDS-PAGE, we noticed that selected proteins seem to be expressed differently in the deletion strains, suggesting that *trmA* and *truB* affect the synthesis of specific proteins.

**Figure 4.**
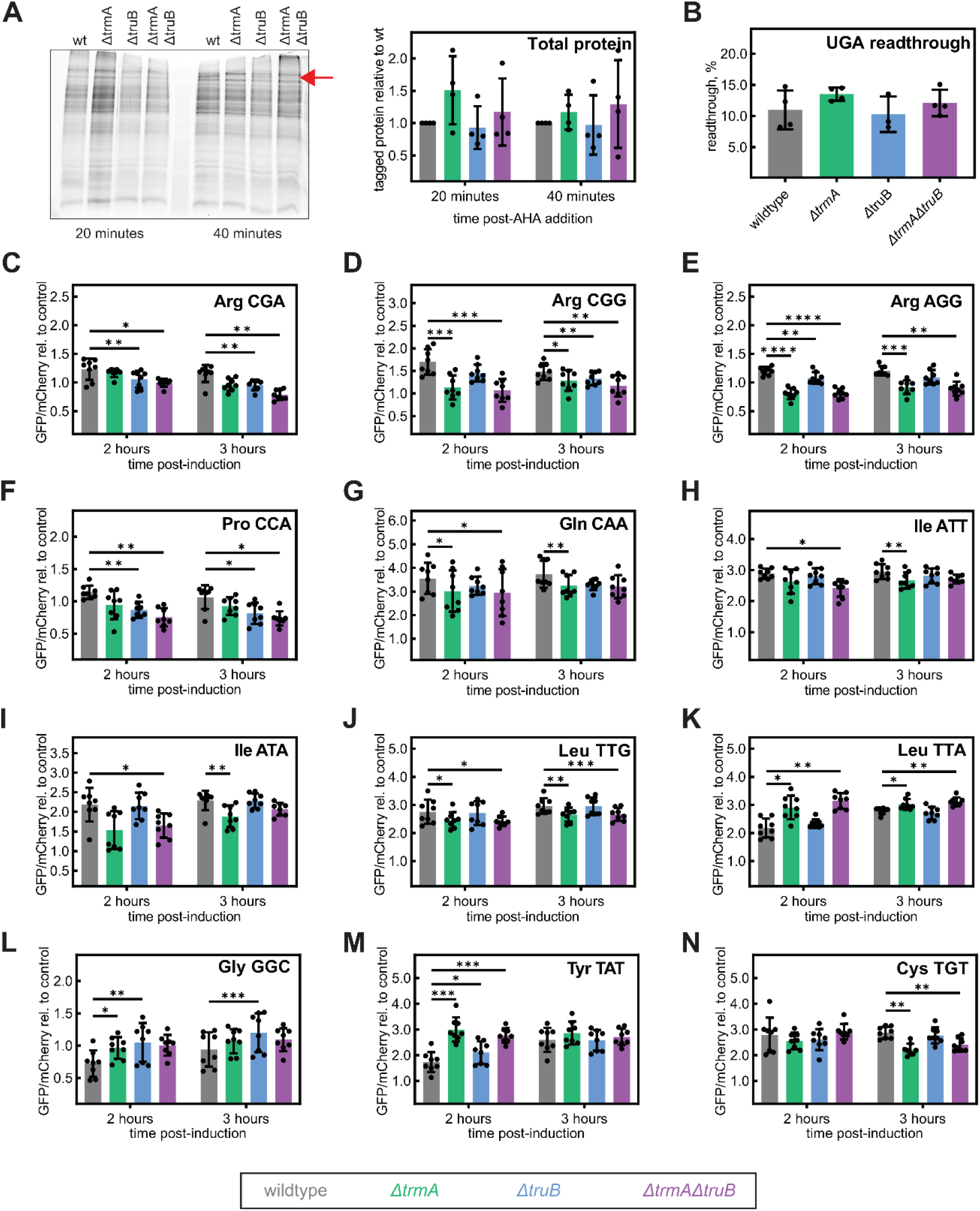
Absence of TrmA and/or TruB does not affect global protein translation, but differentially affects translation of specific codons. **(A)**Determination of global translation by pulse-labeling BONCAT experiments. SDS PAGE shown (left side of panel) is representative of four biological replicates. All replicates are present in Extended Data Figure 4. Red arrow features potential specific protein differences between wildtype and knockout strains. Densitometry of each total lane (right side of panel) was used to estimate global translation in each strain. **(B)** Proportion of UGA readthrough by Sec-tRNA^Sec^ in the context of the *fdhF* SECIS element in a dual luciferase construct. **(C – N)** Measurements of codon-specific translation using GFP reporters. The GFP gene is preceded by four consecutive codons as indicated in the top right for each panel, and GFP expression is recorded 2 and 3 hours after induction relative to mCherry expression encoded on the same plasmid. Codon-specific GFP translation is compared for the *ΔtrmA* (green), *ΔtruB* (blue), *ΔtrmAΔtruB* (purple) and wild-type (grey) *E. coli* strains for indicated codons. Relative translation measurements for the remaining 49 sense codons are present in Extended Data Figure 4 – 6. Eight biological replicates were performed for each strain; * indicates *p* < 0.05, ** indicates *p* < 0.01, and *** indicates *p* < 0.001.

To understand how *trmA* and *truB* affect specific protein synthesis, we analyzed variations in translation for all 61 sense codons in addition to changes in UGA stop codon recoding by the selenoprotein machinery. First, we measured UGA stop codon readthrough by tRNA^Sec^ in response to a selenocysteine insertion sequence (SECIS) element using a luciferase reporter where the UGA codon and the SECIS element of the *E. coli fdhF* gene are placed between a Firefly luciferase and *Renilla* luciferase gene^30^. No significant differences in selonocysteine incorporation are noted between *E. coli* wild-type and the Δ*trmA* and/or Δ*truB* deletion strains (Fig. 4B). Subsequently, we dissected how changes in tRNA maturation and charging affect protein synthesis in a sense codon-specific manner by utilizing a series of reporter constructs where green fluorescent protein (GFP) is preceded by four identical codons ^31^. As an internal reference, mCherry is expressed from the same plasmid. We monitored the ratio of GFP and mCherry fluorescence to determine codon-specific changes in translation in the Δ*trmA* and Δ*truB* single and double-knockout strains (Fig. 4C-N, Extended Data Fig. 4-6). For most codons, we did not observe significant differences between the *E. coli* wild-type and the deletion strains, in accordance with the observation that global translation is unaffected. However, a specific set of codons leads to altered expression of GFP in the Δ*trmA* and/or Δ*truB* deletion strains.

Previously, it was reported that deletion of *truB* (but not *trmA*) results in decreased translation of a reporter containing nine consecutive Arg CGA codons; interestingly, no effect was observed for translation of consecutive CGU codons, which are read by the same tRNA (tRNA^Arg^ACG) ^16^. We detect a similar decrease in translation for only four consecutive CGA codons in strains lacking *truB* (Fig. 4C) and similarly no effect for consecutive Arg CGU codons (Extended Data Fig. 4D). Additionally, we detect translational decreases for other arginine codons, CGG and AGG, which are read by different tRNAs, tRNA^Arg^CGG and tRNA^Arg^CCT, respectively (Fig. 4D, E).

In the Δ*truB* and Δ*trmA*Δ*truB* knockouts, the proline CCA codon (read by tRNA^Pro^TGG) displays significantly lower GFP production (Fig. 4F). Less pronounced, but still significant decreases in translation are noted in the absence of the *trmA* gene for the Gln CAA codon (read by tRNA^Gln^TTG) as well as for the Ile ATT and ATA codons, which are read by tRNA^Ile^GAT and tRNA^Ile2^CAT, respectively (Fig. 4G-I). Interestingly, we observe opposing effects of *trmA* deletion for Leu codon reading: GFP expression is decreased in the presence of consecutive Leu TTG codons (read by tRNA^Leu^CAA), but increased for Leu TTA codons (read by tRNA^Leu^TAA; Fig. 4J, K). For the Gly GGC codon, significantly increased GFP expression is noted in the absence of *truB* (Fig. 4L), and deletion of *trmA* and/or *truB* results in increased translation in the presence of consecutive Tyr TAT codons (Fig. 4M). Knockout of *trmA* decreases translation in the presence of consecutive Cys TGT codons (Fig. 4N). In conclusion, although the translation of many codons is unaffected by *trmA* and/or *truB* translation, certain codons are translated poorly in the absence of *trmA* and/or *truB,* and for other codons translation is enhanced.

### TrmA and TruB impact the E. coli transcriptome and proteome

Since we observed no overall change in protein synthesis, but a change in translation efficiency for individual codons in the absence of *trmA* and/or *truB*, we hypothesized that a subset of proteins, possibly enriched in the codons with altered translation efficiencies, may be differentially synthesized in the deletion strains. To test this hypothesis, we examined RNA abundance in the single- and double-deletion strains by total RNA-seq, and quantitated protein abundance by mass spectrometry, allowing us to correlate the relative abundances of transcripts and proteins in each knockout compared to wildtype (Fig. 5, Extended Data Fig. 7). Overall, these experiments reveal several changes between wild-type and deletion strains on both the transcriptome and proteome level. Particularly, the transcript levels of several genes are increased, but the protein levels are decreased, suggesting deficiencies in translation for the *trmA* and *truB* single and double knockout strains (Fig. 5). This observation is particularly pronounced in the Δ*trmA*Δ*truB* double-knockout strain. Only few proteins increase in abundance despite decreased transcript levels. In general, a larger number of proteins are found to be decreased (rather than increased) upon knockout of *trmA* and/or *truB* compared to wildtype (Extended Data Fig. 7).

**Figure 5.**
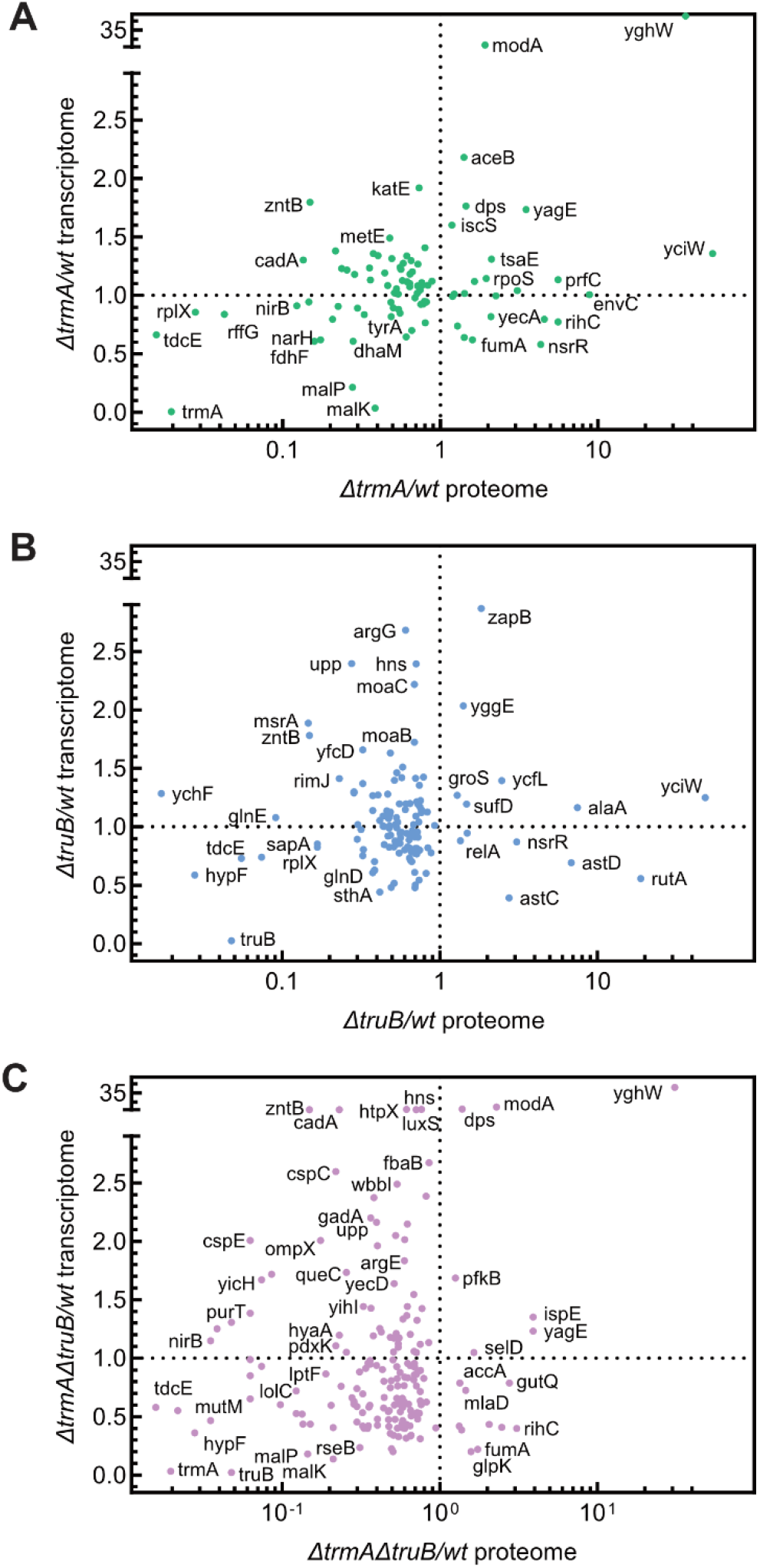
Gene expression is altered upon trmA and/or truB deletion. Transcriptome and proteome changes were recorded in *ΔtrmA* **(A)**, *ΔtruB* **(B)**, and *ΔtrmAΔtruB* **(C)** knockout strains. Each data point represents a protein with a significantly (*p* < 0.05) altered abundance within the knockout compared to wildtype, with fold protein change (knockout/wildtype) indicated by position on the x-axis. To address proteins that were not identified in any wildtype replicates (fold change = division by zero) or not identified in a knockout (fold change = 0), a small value (0.1) was added to each value. For each significantly altered protein, the respective mRNA fold change (knockout/wildtype) is indicated by the value on the y-axis. Proteomics and transcriptomics experiments are the average of four and three biological replicates, respectively.

Analyzing for gene ontology (GO) terms among the down-regulated proteins in the knockout strains (Extended Data Fig. 8) shows that in particular metabolic processes are impacted in the single knockout strains. In the double-knockout strain, both metabolic processes and gene expression processes are affected. While some proteins are decreased in two or three of the knockout strains, other proteins are only decreased in one of the strains (Extended Data Fig. 8) in accordance with strain-specific features observed in the codon-specific GFP translation assays. Overall, these findings are in accordance with our observation that global translation is not affected upon deleting *trmA* or *truB*, but that individual codons are impaired in translation, resulting in specific proteome changes that likely cause the previously reported decrease in cellular fitness^10,11^.

## Discussion

Here, we provide comprehensive insight into the cellular functions of the universally conserved tRNA modifying enzymes TrmA and TruB that introduce m^5^U54 and Ψ55 in all elongator tRNAs (Fig. 6). Thereby, we answer the decades-old question why these enzymes are conserved, but do not display phenotypes upon deletion in many organisms despite their disease relevance in humans and their contributions to cellular fitness. We previously demonstrated that TrmA and TruB fulfill two molecular functions by both modifying and folding tRNAs ^10,11^. To identify their overall function for cellular fitness, we analyzed here the effect of deleting these enzymes in *E. coli* without separating the tRNA modification and folding activities, since our previous data indicate that both are likely contributing to cellular fitness. Specifically, we demonstrate that TrmA and TruB lead to globally increased tRNA aminoacylation without significantly affecting tRNA abundance. Moreover, we uncover functional connections between m^5^U54 and Ψ55 and several other modifications in tRNA (e.g., acp^3^U47, ms^2^i^6^A37, and s^4^U8). In accordance with the absence of major growth phenotypes in Δ*trmA* and/or Δ*truB E. coli* strains, overall translation is unaffected. However, translation for several codons is decreased, most notably the Proline CCA codon and three Arginine codons (CGA, CGG, and AGG). Interestingly, the abundance of several proteins is changed in the Δ*trmA* and/or Δ*truB* strains, which is not correlated with transcript abundance, revealing that the synthesis of specific proteins is impaired. These depleted proteins are often implicated in metabolism or gene expression, which explains the fitness defect of Δ*trmA* and/or Δ*truB* strains. In conclusion, TrmA and TruB enhance aminoacylation and modification of all tRNAs in *E. coli,* thereby fine-tuning their functionality and ensuring efficient protein synthesis of specific codons.

**Figure 6.**
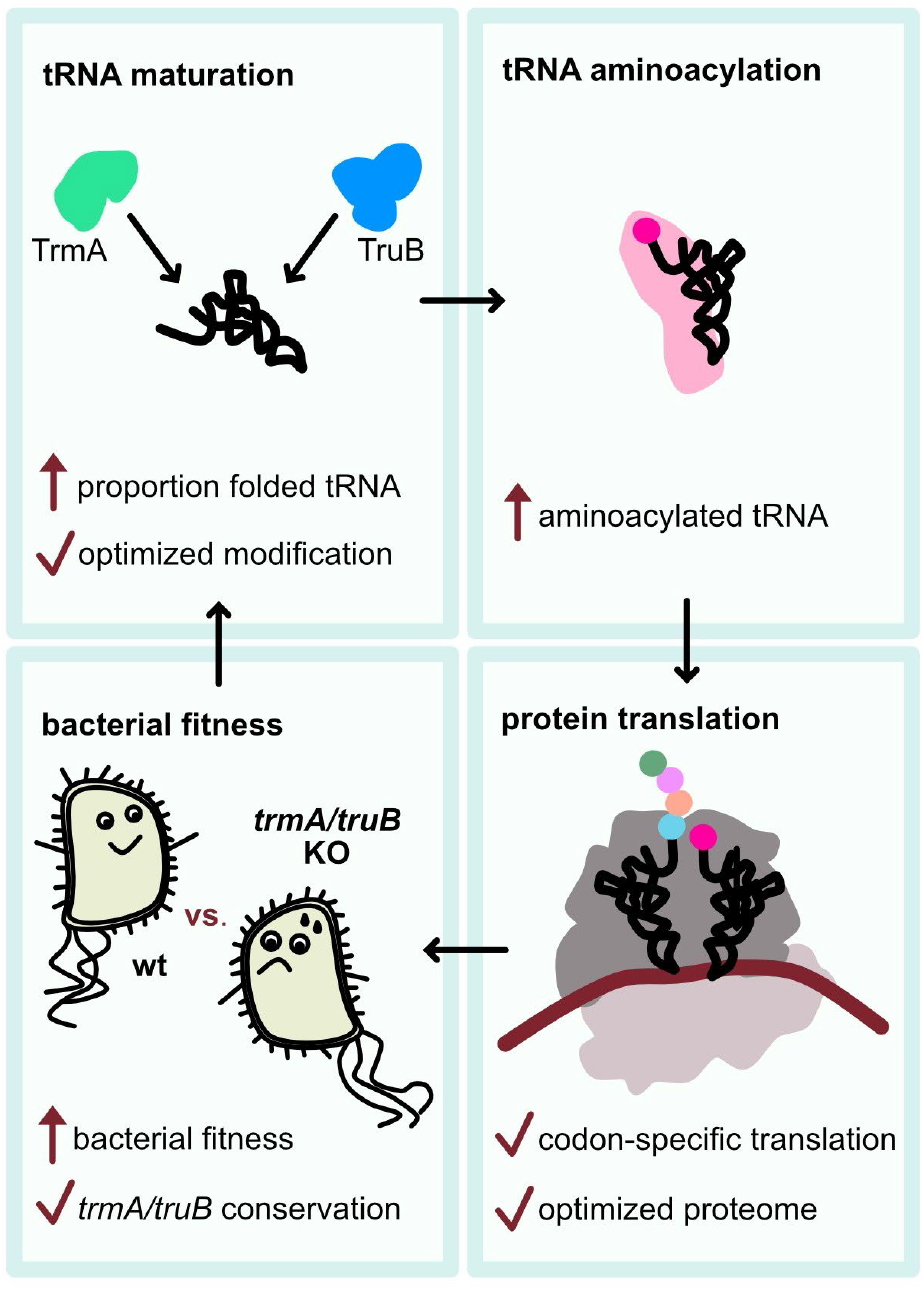
Molecular and cellular function of the tRNA modification enzymes TrmA and TruB. By interacting with the T arm of all tRNA, these enzymes enhance tRNA maturation by introducing m^5^U54 and Ψ55, respectively, promoting tRNA folding, and modulating the formation of several other tRNA modifications. Thereby, they globally increase the levels of tRNA aminoacylation. Whereas these enzymes do not affect overall protein synthesis, translation of several codons is optimized by the presence of TrmA and TruB resulting in a proteome that promotes bacterial fitness in competition assays. The effects of TrmA and TruB on tRNA maturation, aminoacylation, codon-specific translation, the proteome and cellular fitness explain the conservation of these enzymes from bacteria to humans where they are implicated in several diseases.

The data presented here uncover a critical role of TrmA and TruB for tRNA maturation and aminoacylation. Specifically, we reveal global defects in aminoacylation in Δ*trmA* and/or Δ*truB E. coli* strains, wherein presence of TrmA and TruB enzymes collectively increase global aminoacylation by about 10 % compared to the double knockout strain (Fig. 2E-F). Hence, TrmA and TruB affect charging of most or all tRNAs *in vivo,* most likely by enhancing the correct folding of all tRNAs ^10,11^. This decrease in aminoacylation in the knockout strains compared to the wild-type cells is small enough to explain why deletion strains do not display growth defects, but large enough to affect cellular fitness in competition assays over several days ^10,11^. Clearly, it is beneficial for the cell to rescue a small percentage of potentially misfolded tRNAs through the action of the tRNA chaperones TruB and TrmA.

TrmA and TruB and their homologs act early during tRNA maturation, thereby preparing all tRNAs for subsequent maturation events ^32-34^.In both yeast and *Thermus thermophilus,* deletion of *trmA* or *truB* homologs leads to altered abundances of other tRNA modifications at various positions, including m^1^A, Gm, and m^5^U ^14,33^. Here, we reveal that the action of *E.coli* TrmA and TruB modulates the levels of other tRNA modifications including s^4^U8, ms^2^i^6^A37, and acp^3^U47 (Fig. 3). These results agree with recent LC/MS data from the Koutmou lab reporting decreased levels of ms^2^i^6^A37 and acp^3^U47 in tRNA^Phe^ from a *ΔtrmA E. coli* strain (Jones et al.). Together, these findings indicate that many other tRNA modifying enzymes act on tRNA lacking m^5^U54 and/or Ψ55 with reduced efficiency. This effect might be mediated by the chaperone activity of TrmA and TruB, as certain tRNA modification enzymes require correctly folded tRNA, or it may be due to lack of the m^5^U54 and Ψ55 modifications. Further, it is conceivable that the lower levels of other tRNA modifications in the Δ*trmA* and/or Δ*truB* strains contribute to low aminoacylation, as several modifications at positions 34 and 37 act as determinants for aminoacyl-tRNA synthetases ^35^.

Given that m^5^U54 and Ψ55 affect tRNA structural stability ^3^, it is notable that few effects on tRNA abundance were detected in the Δ*trmA* and/or Δ*truB* strains. Only two tRNA^Leu^ isoacceptors are significantly downregulated in the *ΔtrmAΔtruB* strain (Fig. 2C); however, the translation of the Leu CTG and CTA codons read by these tRNAs is not affected by *trmA* and *truB* deletion (Extended Data Fig. 4). This contrasts studies of yeast tRNA modification enzymes, wherein certain enzymes are vital for the stability of a few tRNAs, but dispensable for other tRNAs ^1,26,36,37^. Thus, TruB and TrmA act differently from other tRNA modification enzymes as they do not impair the abundance of selected tRNAs but enhance the maturation of all tRNAs.

Despite the global effect of TrmA and TruB on the maturation and aminoacylation of all tRNAs, there are no global translation changes. However, we observed significant codon-specific defects in translation. In particular, translation of Pro CCA, Arg CGA, CGG and AGG, Gln CAA, Ile ATT and ATA, Cys TGT, and Leu codons TTG were negatively affected by the deletion of the *trmA* and/or *truB* gene. These codons are all used at much lower frequencies in *E. coli* than the other codons for the same amino acid. Accordingly, the tRNAs decoding these codons are often at a lower abundance compared to their isoacceptors; however, the decrease in their activity cannot be attributed to a reduction in their abundance compared to the knockout strains (Fig. 2A-C). Instead, these tRNAs are particularly susceptible to defects in codon-specific decoding abilities in the absence of TrmA and TruB, whether that be due to the loss TrmA/TruB chaperone activity and/or the loss of the m^5^U54 and Ψ55 modifications. Alternatively, defects in these tRNAs may caused by changes in the activity of other tRNA modifying enzymes, including those that form ms^2^i^6^A37 or acp^3^U47, as these modifications tend to be decreased within the tRNAs affected in our codon-specific translation assay.

Of the seven tRNAs that contain acp^3^U47, six have decreased acp^3^U47 levels in the *ΔtrmAΔtruB* knockout strain. Remarkedly, these six tRNAs decode at least one codon differently in the absence of *trmA* (and *truB*) compared to the wildtype strain (Fig. 3C). Translation of CGA by tRNA^Arg^ACG, ATT and ATA by tRNA^Ile^GAT, ATA by tRNA^Ile2^CAT, AAA by tRNA^Lys^TTT, and GTT by tRNA^Val^GAC are significantly lower in the *ΔtrmA* and/or *ΔtruB* strain, whereas translation of Phe TTC codons is improved in the *ΔtrmA* strain. The ms^2^i^6^A37 modification also has a lower abundance in several tRNAs within the *ΔtrmA* strain (Fig 3F), and half (4/8) of the ms^2^i^6^A37-containing tRNAs display different translation of certain codons in the absence of *trmA*. In the absence of *trmA* (and *truB*), translation is decreased for TGT and TTG, read by tRNA^Cys^GCA or tRNA^Leu^CAA and tRNA^Leu^TAA, respectively. In contrast, translation is increased for TAT read by tRNA^Tyr^GTA and TTC read by tRNA^Phe^GAA. Notably, several hypomodified tRNAs discussed above also decode additional codons whose translation is not altered in the absence of *trmA/truB*, supporting the notion that TrmA and TruB modulate codon-specific tRNA interactions. Interestingly, recent *in vitro* translation assays by the Koutmou group suggest translocation in the presence of the antibiotic hygromycin B is improved in the absence of m^5^U54 (Jones et al.), providing further evidence for tRNA elbow modifications altering translation. The observed tRNA- and codon-specific effects in the absence of m^5^U54 and Ψ55 could be explained by changes in the tRNA dynamics which are critical when the tRNAs translocated through the ribosome.

In accordance with codon-specific translational impairments in the knockout strains, we noted a decreased abundance of specific proteins in the Δ*trmA* and/or Δ*truB* strains although their transcripts levels were unchanged or even increased, demonstrating translation-specific defects in gene expression. Indeed, analysis of the most decreased proteins in the Δ*trmA*Δ*truB* strain shows that these are often enriched in the most affected codons such as the CGA and CGG Arg codons (ZntB) and the CCA Pro codon (YicH, PurT, NirB). There are 20 proteins whose abundance is decreased in all three single- and double-knockout strains (Extended Data Fig. 8); nine genes encoding these proteins are enriched in Pro CCA and Arg CGA and CGG codons (*adhP*, *hrpA*, *hyaA*, *rbsK*, *upp*, *ybiS*, *ycaO*, *ygaM*, and zntB). Evidently, the codon-specific defects in translation upon *trmA* and/or *truB* deletion affect the overall proteome underlining the importance of these two tRNA modifying enzymes for protein synthesis. Moreover, 9 of the 20 translationally downregulated proteins are large *E. coli* proteins with more than 500 residues (*rpoC*, *fusA*, *hrpA*, *hyaB*, *metE*, *nrdA*, *pheT*, *tdcE*, *ycaO*). For these large proteins, subtle effects on translation in the Δ*trmA* and/or Δ*truB* strains likely accumulate leading to lower protein abundance. The codon-specific translation effects and decreases in selected proteins explain the reduced cellular fitness of the Δ*trmA* and/or Δ*truB* strains.

As TrmA and TruB are conserved across organisms from bacteria to humans, the link between their molecular function during tRNA maturation and aminoacylation and their cellular role for protein synthesis is likely similar in eukaryotes. In yeast, studies similarly suggest a role for the TrmA and TruB homologs, Trm2 and Pus4, in translation. Global ribosome profiling has revealed both Trm2 and Pus4 have subtle effects on ribosome occupancy at different codons, suggesting codon-specific translation defects in yeast knockout strains, similar to our observations in bacteria ^38^. Moreover, yeast Pus4 can act as a prion, and protein synthesis is altered in these prion strains likely due to a different translation of rare codons ^39^. The human homologs, TRMT2A and TRUB1 have also been implicated in modulating protein synthesis. Although human TRMT2A does not affect synthesis of polyQ proteins, the aggregation of polyQ is reduced ^19^ and translation fidelity is decreased in the absence of TRMT2A ^40^. Moreover, human TRUB1 affects mitochondrial tRNA conformation and mitochondrial translation thus decreasing the assembly and activity of oxidative phosphorylation complexes, without altering tRNA abundance ^21^. In conclusion, our findings regarding the molecular and cellular function of *E. coli* TrmA and TruB, together with reports of the importance of their yeast and human homologs for protein synthesis indicates a conserved mechanism for these tRNA modifying enzymes. Herein, TrmA and TruB homologs promote tRNA maturation and aminoacylation and affect codon-specific translation thus providing an explanation for their implication in human diseases, including diabetes, deafness, and cancer ^18-21^.

In summary, we comprehensively identify the functions of the conserved tRNA modifying enzymes, TrmA and TruB, which introduce m^5^U54 and Ψ55 in the TΨC arm of all tRNAs (Fig. 6). Thereby, these enzymes promote correct tRNA folding and enable efficient modification by other tRNA modifying enzymes. These positive effects during maturation lead to significant improvements in aminoacylation for all tRNAs. Thus, TruB and TrmA stand out as enzymes that globally affect all tRNAs, contrasting other tRNA modifying enzymes which only act on a subset of tRNAs ^26^. Other tRNA modifying enzymes targeting the elbow region of many tRNAs such as the dihydrouridine synthases may have similar global roles. The positive impact of TrmA and TruB on the cell is particularly evident in the decoding of codons by tRNAs harboring ms^2^i^6^A37 and acp^3^U47 modifications and the translation of large proteins leading to significant changes in protein expression in Δ*trmA* and/or Δ*truB*knockout strains. These codon- and protein-specific translation defects reduce the cellular fitness under competition conditions. Similarly, the human homologs of *E. coli* TrmA and TruB are likely to impact tRNA maturation and aminoacylation and to modulate protein synthesis explaining their implication in human diseases. Unlike other tRNA modification enzymes that have tRNA- and stress-specific functions, we therefore demonstrate that TrmA and TruB act as global enhancers of protein synthesis and cellular fitness.

## Methods

### Chemicals and reagents

Unless otherwise stated, all reagents were purchased from ThermoFisher Scientific. All oligonucleotides were purchased from Integrated DNA Technologies (IDT) and are listed in Table S1.

### E. coli strains

All *E. coli* wildtype and single knockout strains are BW25113 derivatives from the Keio Collection ^22^. To construct the *ΔtrmAΔtruB* double knockout, the FLP recognition target (FRT)-flanked kanamycin resistance cassette was removed from the Keio *ΔtrmA* strain, and the *truB* gene was subsequently replaced by a FRT-flanked kanamycin resistance cassette using λ Red recombinase ^41^. All gene deletions were confirmed by colony PCR using primers specific for the kanamycin resistance cassette (k1/k2) and primers targeting the locus upstream and downstream of the deleted gene (Table S1).

### Total RNA extractions for tRNAseq, Northern blotting, and primer extension

Cells were grown overnight in LB medium (wildtype) including 50 µg/mL kanamycin (knockout strains) from single colonies. Cultures were centrifuged and resuspended in fresh LB media or LB including 2% (w/v) sodium formate as indicated. Cultures (50 mL) were started at an OD600 of 0.1 and grown until reaching an OD600 of 0.3-0.4, when 15 mL culture was harvested and shock frozen. RNA extractions were accomplished under acidic conditions (pH <5) to preserve the aminoacyl bond using TRIzol (Invitrogen), following the manufacturer’s instructions. In brief, cells were resuspended in 3 mL TRIzol, followed by the addition of 0.6 mL chloroform. After centrifugation (30 minutes, 6000 *g*), the aqueous layer was removed and ethanol precipitated several times. Purified RNA was resuspended in 10 mM NaOAc (pH 4.8) and stored at -80 °C. Concentrations were determined using UV spectrometry (NanoDrop 2000c).

### MSR-tRNA-seq for tRNA abundance, aminoacylation, and certain modifications

Library preparation, MSR-tRNA-seq and data analysis were conducted as previously published ^24^. Three biological replicates were sequenced for each strain and condition. The *E. coli* tRNA reference was obtained from Genomic tRNA database ^42^, Modification was identified by mutation and deletion signatures that were generated by Superscript IV read-through. Source codes for all custom scripts are available on GitHub (https://github.com/ckatanski/CHRIS-seq).

### Northern blotting for tRNA thiolation

To compare the fraction of thiolated tRNAs to non-thiolated tRNAs, 5 µg total RNA was separated on a 20 µM [(N-acryloylamino)phenyl]mercuric chloride (APM, Toronto Research Chemicals) 7 M urea 12% polyacrylamide gel ^28^. To reduce APM-thiol linkages, APM gels were soaked in 0.2 M β-mercaptoethanol for one hour prior to transfer. RNA was transferred to a nylon membrane (Amersham Hybond-XL) at 23 V for 7 minutes using a Trans-Blot Turbo Transfer System (Bio-Rad). Following transfer, blots were crosslinked twice at 1200 mJ. Northern blotting was performed similarly as described in ^43^ using biotinylated probes (Table S1), which were detected after incubation with 120 ng/mL streptavidin-HRP (Genscript) in hybridization buffer for one hour. To detect chemiluminescence, a solution containing 375 ng/mL luminol, 55 ng/mL p-coumaric acid, and 0.1% (v/v) hydrogen peroxide was applied to the blot immediately prior to imaging with a FluorChem Q imager (Cell Biosciences). Densitometry was performed using ImageJ.

### Primer extension to detect acp^3^U47 modification

Reverse transcriptions were performed by first annealing 1700 ng of total RNA to 3 pmol Cy5-labeled tRNA^Phe^ reverse transcription primer by heating to 65 °C and cooling to 47 °C. The RNA-primer mixture was then added to a reaction mixture containing final concentrations of 136 ng/µL total RNA, 240 nM Cy5-labeled reverse transcription primer, 0.4 mM dNTPs, and 8 mM DTT, in 1 X AMV reverse transcription buffer. After pre-heating to 47 °C, AMV reverse transcriptase (New England Biolabs) was added to a final concentration of 0.4 U/µL. and incubated at 47 °C for 45 minutes. The reaction was ended by heating to 70 °C for 15 minutes. Samples (7 µL of the reverse transcription reaction) were analyzed on an 8 M urea 8% polyacrylamide sequencing gel in TBE which was preheated to 50 °C at 50 W until the bromophenol blue dye front was halfway down the gel. Gels were imaged on a Typhoon 5 scanner.

### Total RNA sequencing

As TRIzol extractions from *E. coli* cells are known to be biased towards the extraction of small RNAs ^44^, and to promote proportionate presence of mRNAs of all sizes, *E. coli* cells were first resuspended in 300 µL of a cell opening buffer (100 mM Tris, 10 mM EDTA, 0.5% SDS, 286 mM β-mercaptoethanol) containing 10 mg/mL lysozyme, 0.1 U/µL DNase I, and 0.1 U/µL RiboLock RNase inhibitor (ThermoFisher) and incubated at 37 °C for 5 minutes. Subsequently, Proteinase K (ThermoFisher) was added to a final concentration of 0.5 mg/mL, and lysates were incubated at 50 °C for 10 minutes. RNA extractions were performed as described above. Further DNase treatment, RNA quality analysis by TapeStation and DNA/RNA Qubit, library preparation, strand-specific Illumina sequencing with a HiSeq400, and data analysis were performed by Azenta. In brief, reads were trimmed with Trimmomatic v.0.36 and trimmed reads were mapped to the *E. coli* BW25113 genome using Bowtie2 v.2.2.6. Unique gene hit counts were calculated by using featureCounts from the Subread package v.1.5.2 and gene expression was compared between *E. coli* strains using DESeq2.

### Bioorthogonal noncanonical amino acid tagging (BONCAT)

*E. coli* pre-cultures were grown overnight in the appropriate antibiotic in 5 mL LB medium. Stationary-phase cells were harvested, washed, and resuspended in M9 minimal media containing 1.75% sodium formate, and 50 mL cultures were grown at 37°C starting at an OD600 of 0.1. When cultures reached an OD600 of ∼0.3, L-azidohomoalanine (L-AHA, Sigma) was added to a final concentration of 1 mM. Twenty minutes and 40 minutes post-L-AHA addition, 0.5 OD600 units of cells were harvested, fixed with 3% (v/v) formaldehyde, washed into PBS, and stored at -20 °C. Cell opening and fluorescent labeling of L-AHA-containing proteins with DBCO-PEG4-Carboxyrhodamine 110 (Click Chemistry Tools) were performed as described ^29^. Proteins were purified using chloroform-methanol extraction and resuspended in 6.4 M urea and 200 mM DTT; total protein concentration was determined by UV spectrometry:

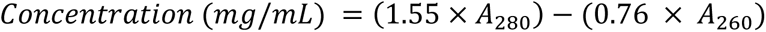

Protein samples were denatured at 65 °C prior to analyzing 5 µg total protein on a 12% SDS-PAGE. Labeled proteins were detected using the 488 nm laser on a Typhoon 5 Imager (Cytiva). To confirm equal loading, gels were stained with Coomassie. Relative global translation for each strain was estimated by densitometry (ImageJ).

### Proteome comparison by mass spectrometry

Four biological replicates of each strain were grown at 37 °C in LB media containing 2% (w/v) sodium formate until reaching an OD600 between 0.3 and 0.4. Following harvesting, cells were resuspended in 8 M urea containing SDS loading buffer lacking any colored dyes and 0.3 OD600 units of each sample were analyzed on a 12% SDS PAGE and stained with Coomassie.

Each lane was excised and subsequently cut into 14 equal bands, each gel fraction was subjected to in-gel tryptic digestion as previously described ^45^ and the resulting peptides were dried and resuspended in 60 L of 0.2% formic acid in 5% acetonitrile (ACN). Digested peptides were analyzed by LC-MS/MS using a ThermoScientific Easy nLC-1000 in tandem with a Q-Exactive Orbitrap mass spectrometer. Each sample (5 uL) was resolved using a 120 min gradient [0%–45% Buffer B; Buffer A (0.2% formic acid in 5% ACN); Buffer B (0.2% formic acid in 100% ACN)] on a 2 cm Acclaim 100 PepMap Nanoviper C18 trapping column in tandem with a Thermo EASY-Spray column (PepMap® RSLC, C18, 3 um, 100 Å, 75 m × 150 mm). For data-dependent analysis, full scans were acquired at 35000 resolution at a range of 400–200 m/z while 17500 resolution was used for MS/MS scans. Only top 15 ions with +2 and +3 charges were selected for MS/MS with 10 s dynamic exclusion applied to prevent continuous reanalysis of abundant peptides. Following data acquisition, raw data files were compiled for each gel lane and searched with Proteome Discoverer 1.4’s SEQUEST search algorithm using the reviewed, non-redundant *E. coli* complete proteome retrieved from UniProtKB. The search parameters and quantification were as previously described ^46^. Proteins were considered to be significantly different in each knockout compared to the wildtype for p values <0.05 determined by a two-tailed paired Student’s t test. Significantly enriched GO terms in each dataset were determined using PANTHER GO biological complete annotation set and *E. coli* reference list by Fisher’s Exact test, correcting for False Discovery Rate ^47^.

### Translation reporter assays to examine codon-specific translation

BW25113 wildtype, *ΔtrmA*, *ΔtruB,* and *ΔtrmAΔtruB* strains were transformed with reporter plasmids carrying an arabinose-inducible fluorescent transcriptional cassette encoding superfolder green fluorescent protein (sfGFP) followed by mCherry fluorescent protein (kind gift of Assaf Katz, Universidad de Chile, Santiago, Chile) ^31^. Each reporter contains a set of four tandem repeats of a codon following the third sfGFP codon, such that sfGFP expression relies on translation of the four repeated codons. To account for small differences in translation between strains, plasmid S1 was used as a control, which lacks any additional codons in sfGFP.

sfGFP and mCherry fluorescence was determined similarly as previously described ^31^. In short, overnight cultures were diluted 1:20 to an OD600 of ∼0.1 and grown in LB medium supplemented with 100 µg/mL ampicillin until reaching an OD600 of 0.4-0.6. At this point, cells were diluted 1:4 into fresh LB medium containing a final concentration of 100 µg/mL ampicillin and 0.4% (w/v) arabinose in black, optical bottom 96 well plates and grown at 37 °C with shaking at 100 rpm. Two- and three-hours post-induction, OD600, sfGFP fluorescence (excitation: 480 nm, emission: 515 nm), and mCherry fluorescence (excitation: 587 nm, emission: 610 nm) were measured using a FlexStation 3 plate reader (Molecular Devices). sfGFP/mCherry ratios for each test codon were normalized to the sfGFP/mCherry ratio of plasmid S1 for the respective strain:

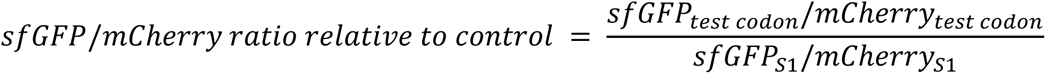

For each codon, translation between strains was compared using two-way ANOVA. Significant differences (p <0.05) between strains are indicated.

### Sec-tRNA^Sec^ mediated UGA readthrough reporter assay

To prepare a dual luciferase reporter for measuring selenocysteine insertion sequence-directed UGA readthrough that is suitable for expression within kanamycin-resistant and DE3-lacking BW25113 knockout strains, pTrc99a-fluc-*fdhF*130-179UGA-rluc was prepared by PCR amplification and ligation of the DNA sequence encoding the Fluc-FdhF-Rluc fusion protein within pET24-fluc-fdhF130-179UGA-rluc ^30^ (kind gift from Marina Rodnina, Max Planck Institute for Multidisciplinary Sciences, Göttingen, Germany). To account for subtle differences in canonical translation between strains, site-directed mutagenesis was used to construct the control plasmid, pTrc99a-fluc-*fdhF*130-179UUC-rluc, which encodes a UUC Phe codon in place of the UGA Sec codon. Plasmids were transformed into BW25113 wildtype, *ΔtrmA*, *ΔtruB,* and *ΔtrmAΔtruB* strains.

To determine the proportion of UGA readthrough in each strain, cultures were grown at 37 °C with shaking, starting at an OD600 of ∼0.1 in LB media supplemented with 50 µM sodium selenite. At an OD600 between 0.5-0.7, fusion protein expression was induced with 1 mM isopropyl ß-D-1-thiogalactopyranoside (IPTG). Following 30 minutes of growth, 0.1 OD600 of cells were harvested, opened by lysozyme, and firefly and *Renilla* luciferase were measured using the Dual-Luciferase® Reporter Assay System (Promega) as per the manufacturer’s instructions with a Spectramax i3 plate reader (Molecular Devices). Recoding efficiency for each strain was calculated using the following formula:

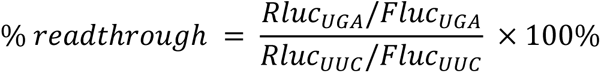

### Submission numbers

The data discussed in his publication have been deposited in NCBI’s Gene Expression Omnibus and are accessible through GEO Series accession number GSE233667 for MSR-tRNA-seq data and GSE237609 for total RNA sequencing data.

## Acknowledgements

We thank Assaf Katz for sharing a complete set of codon-specific reporter assays and Marina Rodnina for sharing a reporter construct for selenocysteine incorporation during translation. Anileen Pageni, Kieran Meadows, Kevin Li and Timothy Vos have helped with the preparation and initial characterization of *E. coli* strains which is greatly appreciated. This work was supported by the Natural Sciences and Engineering Research Council of Canada [UK: Discovery Grant RGPIN-2020-04965 and Discovery Accelerator Supplement RGPAS-2020-00010].

## Extended Data

**Extended Data Figure 1.**
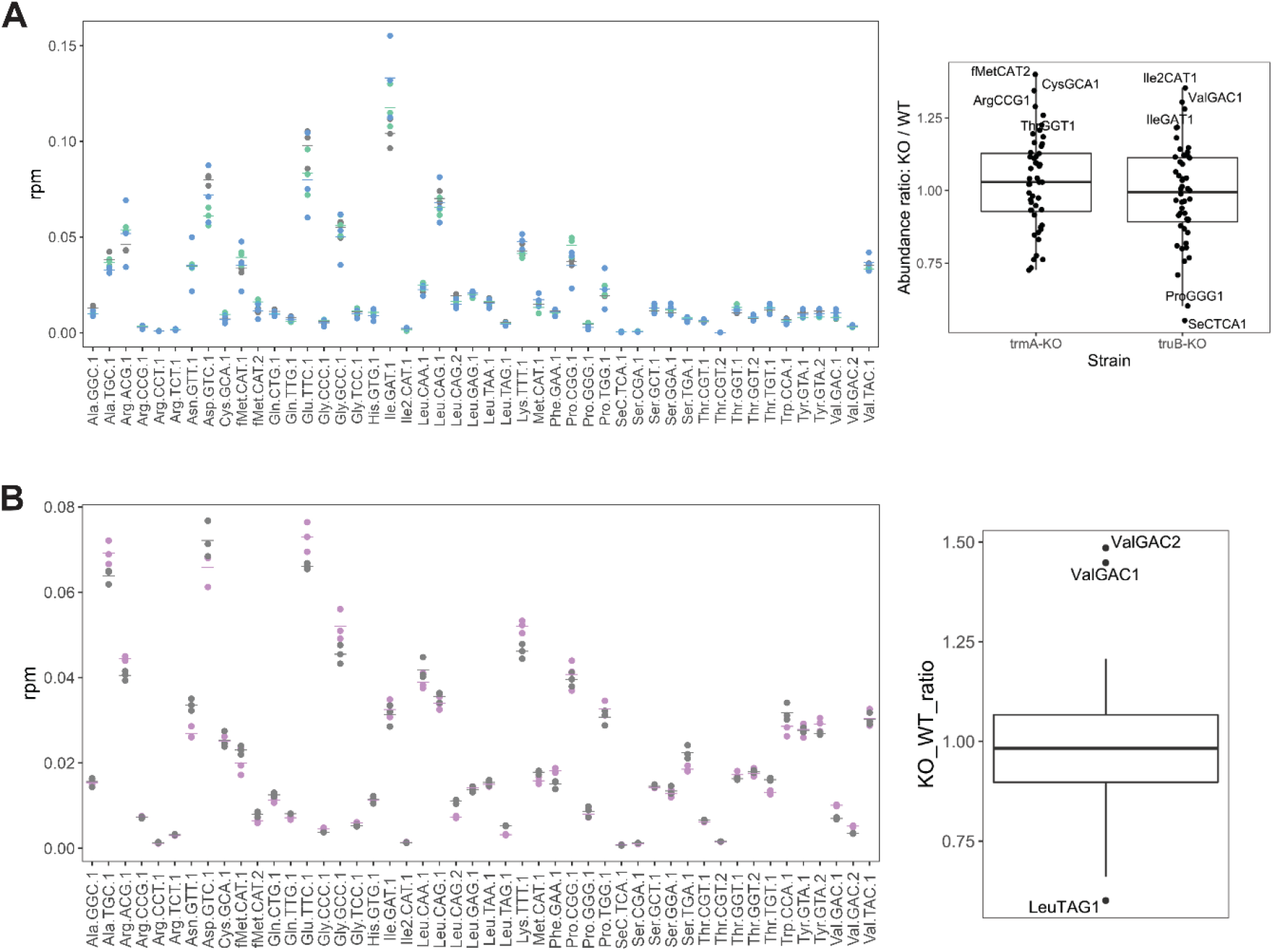
Global tRNA abundance does not change in the absence of *trmA* and/or *truB* observed by RNAseq. Reads per million (RPM) counted for each tRNA species in wildtype and single *ΔtrmA* and *ΔtruB* knockout (KO) strains **(A)** or wildtype and the double *ΔtrmAΔtruB* strain **(B)**.

**Extended Data Figure 2.**
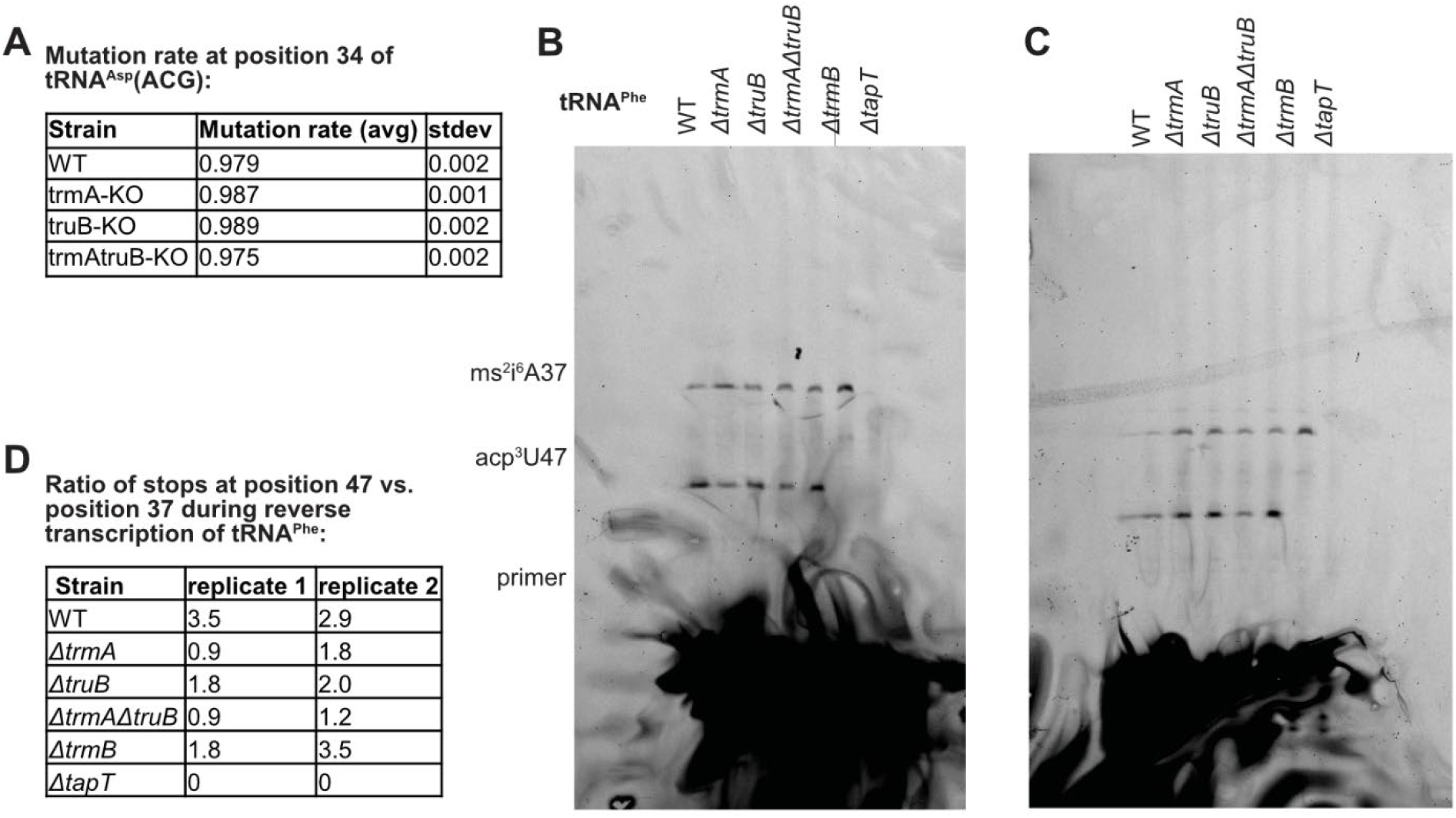
tRNA modifications in the absence of *trmA* and/or *truB*. **(A)**Mutation rate observed for position 34 of tRNA^Arg^ACG, which indicates presence of the inosine modification. **(B)(C)** Full, unedited gel images for two biological replicates (replicate 1, C; replicate 2, D) for tRNA^Phe^ primer extension to detect acp^3^U47 modification. To determine the relative acp^3^U47 content between strains, the reverse transcription stop corresponding to acp^3^U47 was compared to that for ms^2^i^6^A37 **(D).**

**Extended Data Figure 3.**
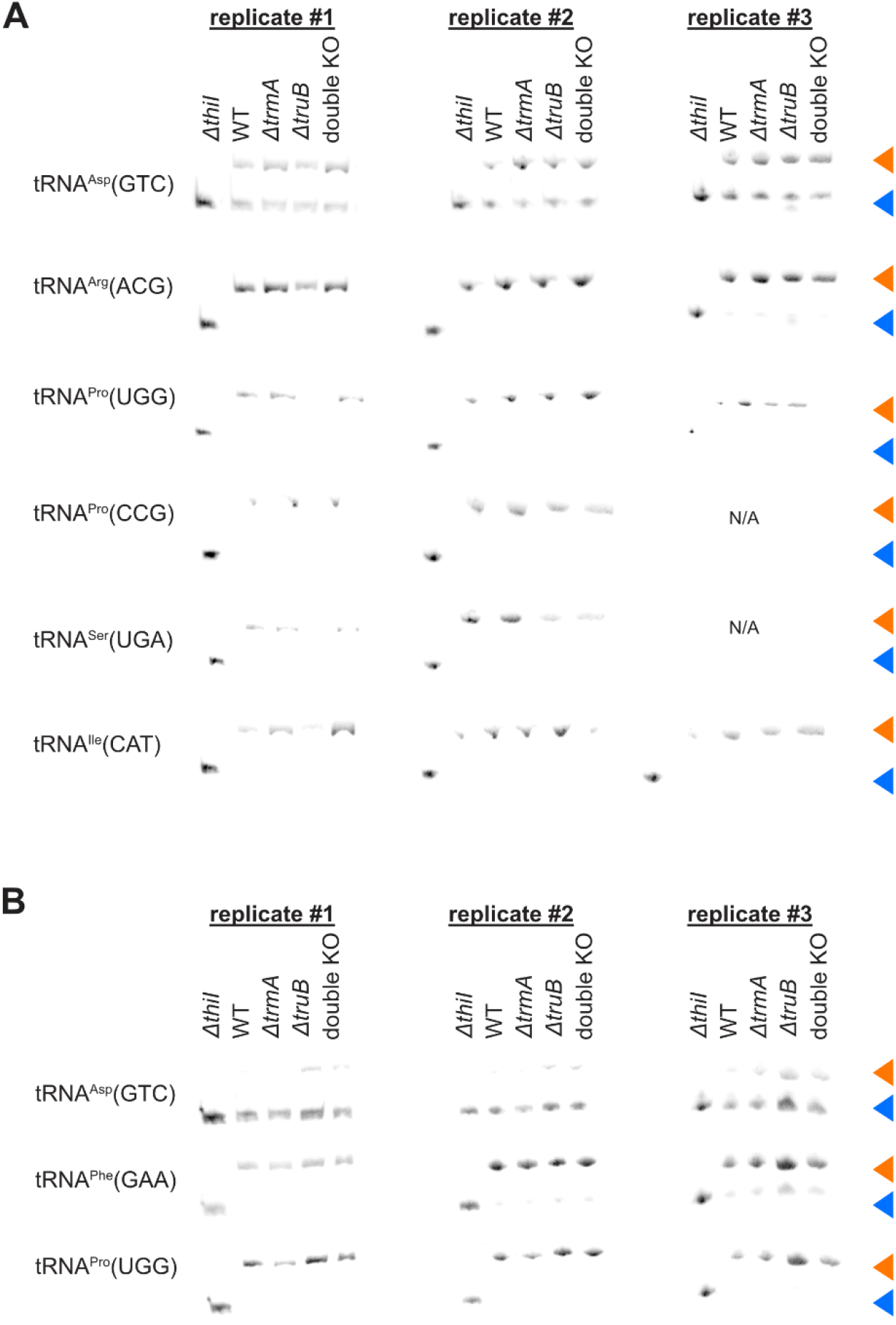
APM Northern blotting to determine the levels of s^4^U8 in different tRNAs. **(A)***E. coli* strains were grown in rich LB media, total tRNA was extracted and thiolation levels of specific tRNAs were determined by APM-Urea PAGE followed by detection using northern blotting. No significant differences between tRNA thiolation for these tRNAs were identified in this growth condition. **(B)** Thiolation levels of specific tRNAs were determined similar as in (A) after growing *E. coli* under stress conditions in LB media containing 2% sodium formate. The thiolated tRNA migrates slower and is indicated with an orange arrow in contrast to the non-thiolated form (blue triangle). Thiolation levels were found to be different in *trmA* and *truB* knockout strains compared to wildtype for tRNA^Asp^GTC (see Figure 3).

**Extended Data Figure 4.**
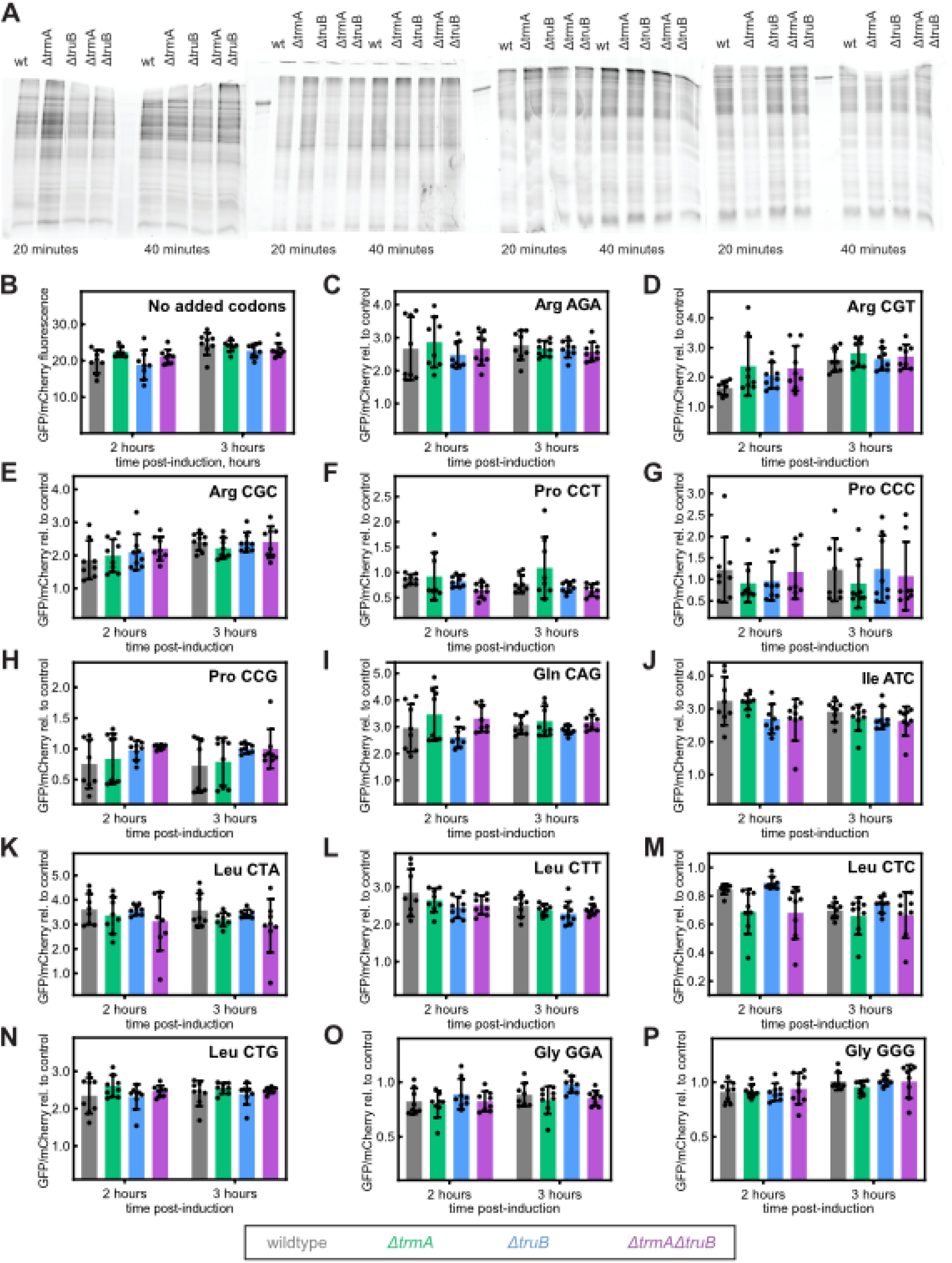
Absence of TrmA and/or TruB disproportionately affects translation of specific codons (set 1). **(A)**Original SDS-PAGEs detecting newly synthesized proteins in the *ΔtrmA*, *ΔtruB*, *ΔtrmAΔtruB* and wild-type *E. coli* strains. **(B to P)** The GFP reporter gene is preceded by four consecutive codons as indicated in the top right for each panel, and GFP expression is compared 2 and 3 hours after induction relative to mCherry expression encoded on the same plasmid. Codon-specific GFP translation is reported for the *ΔtrmA* (green), *ΔtruB* (blue), *ΔtrmAΔtruB* (purple) and wild-type (grey) *E. coli* strains. Significant changes are marked with an asterisk. In panel (B), a control is shown where no consecutive codons precede the GFP gene showing that general translation is not affected in the knockout strains.

**Extended Data Figure 5.**
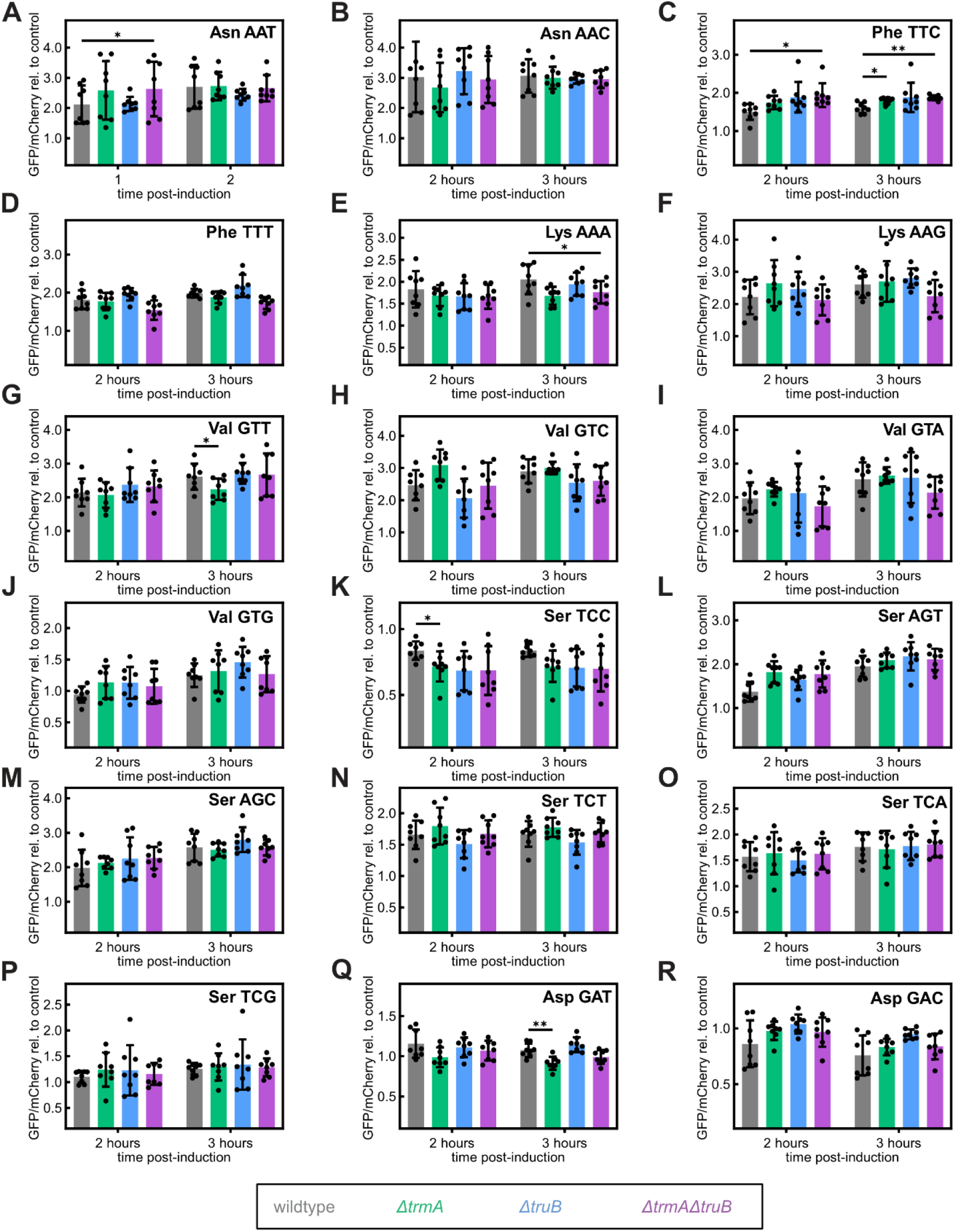
Absence of TrmA and/or TruB disproportionately affects translation of specific codons (set 2). The GFP reporter gene is preceded by four consecutive codons as indicated in the top right for each panel, and GFP expression is compared 2 and 3 hours after induction relative to mCherry expression encoded on the same plasmid. Codon-specific GFP translation is reported for the *ΔtrmA* (green), *ΔtruB* (blue), *ΔtrmAΔtruB* (purple) and wild-type (grey) *E. coli* strains. Significant changes (*p* < 0.05) are marked with an asterisk.

**Extended Data Figure 6.**
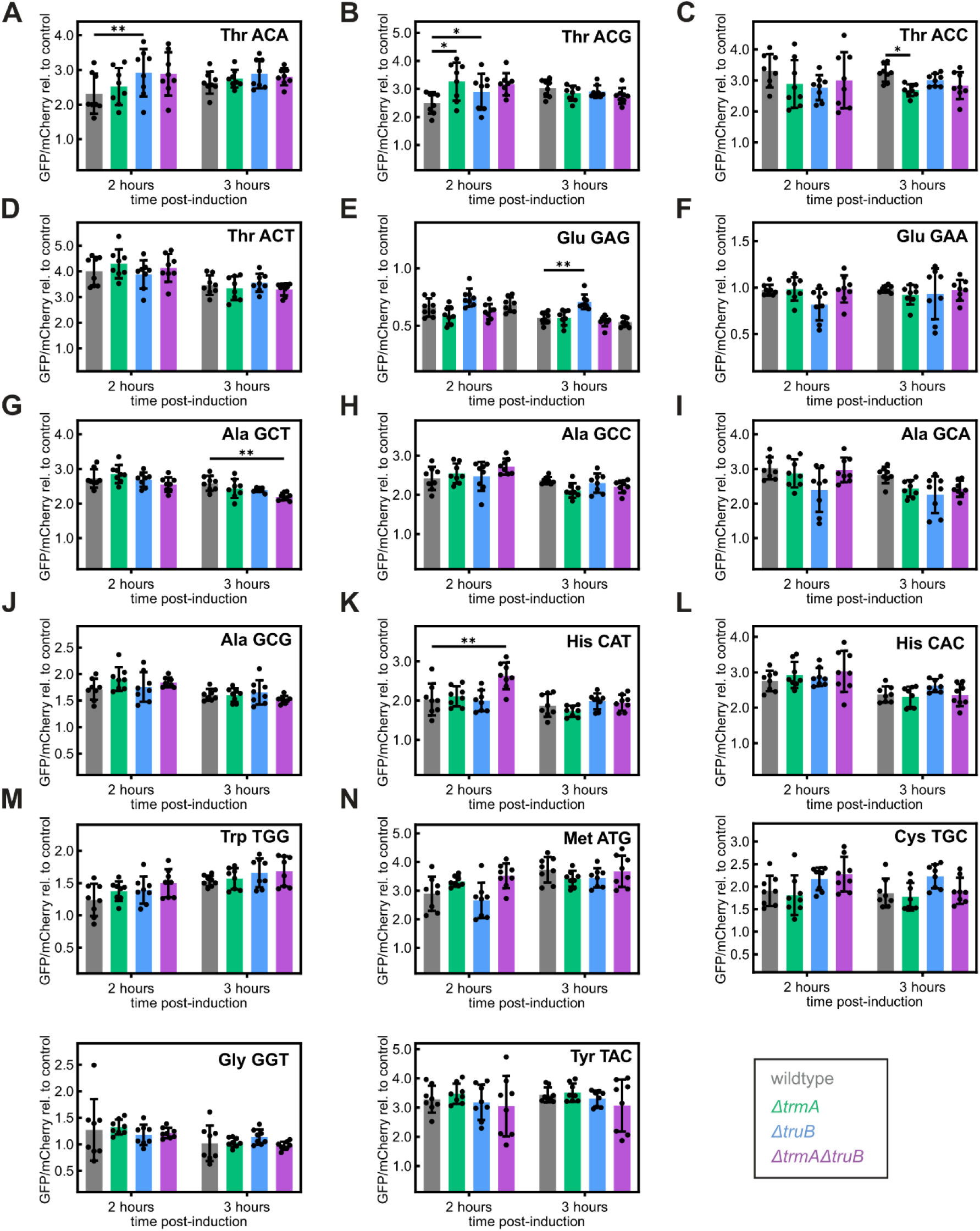
Absence of TrmA and/or TruB disproportionately affects translation of specific codons (set 3). The GFP reporter gene is preceded by four consecutive codons as indicated in the top right for each panel, and GFP expression is compared 2 and 3 hours after induction relative to mCherry expression encoded on the same plasmid. Codon-specific GFP translation is reported for the *ΔtrmA* (green), *ΔtruB* (blue), *ΔtrmAΔtruB* (purple) and wild-type (grey) *E. coli* strains. Significant changes (*p* < 0.05) are marked with an asterisk.

**Extended Data Figure 7.**
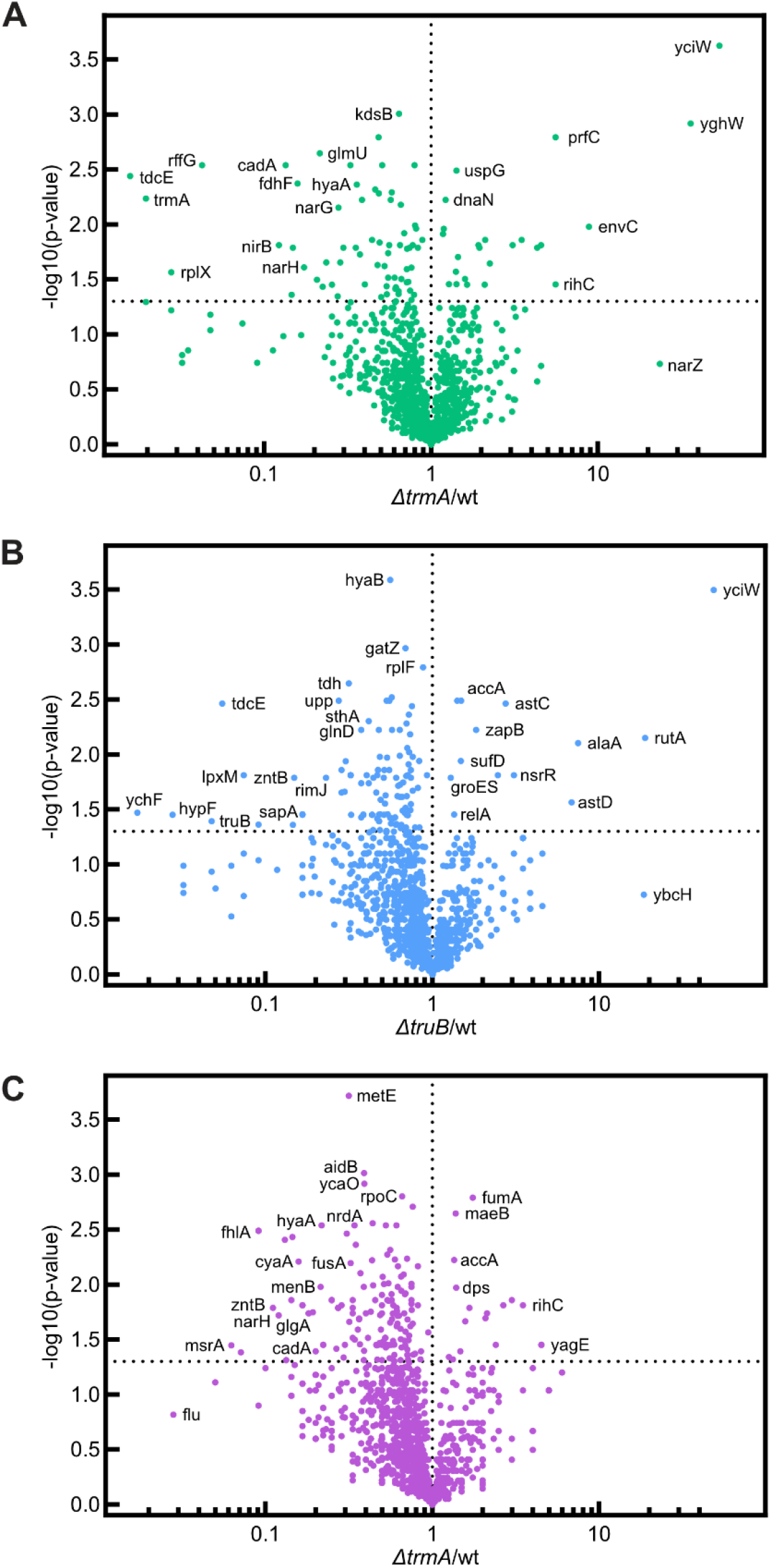
Volcano plots showing the change in relative protein abundance between E. coli wildtype and *ΔtrmA* (A), *ΔtruB* (B), and *ΔtrmAΔtruB* (C).

**Extended Data Figure 8.**
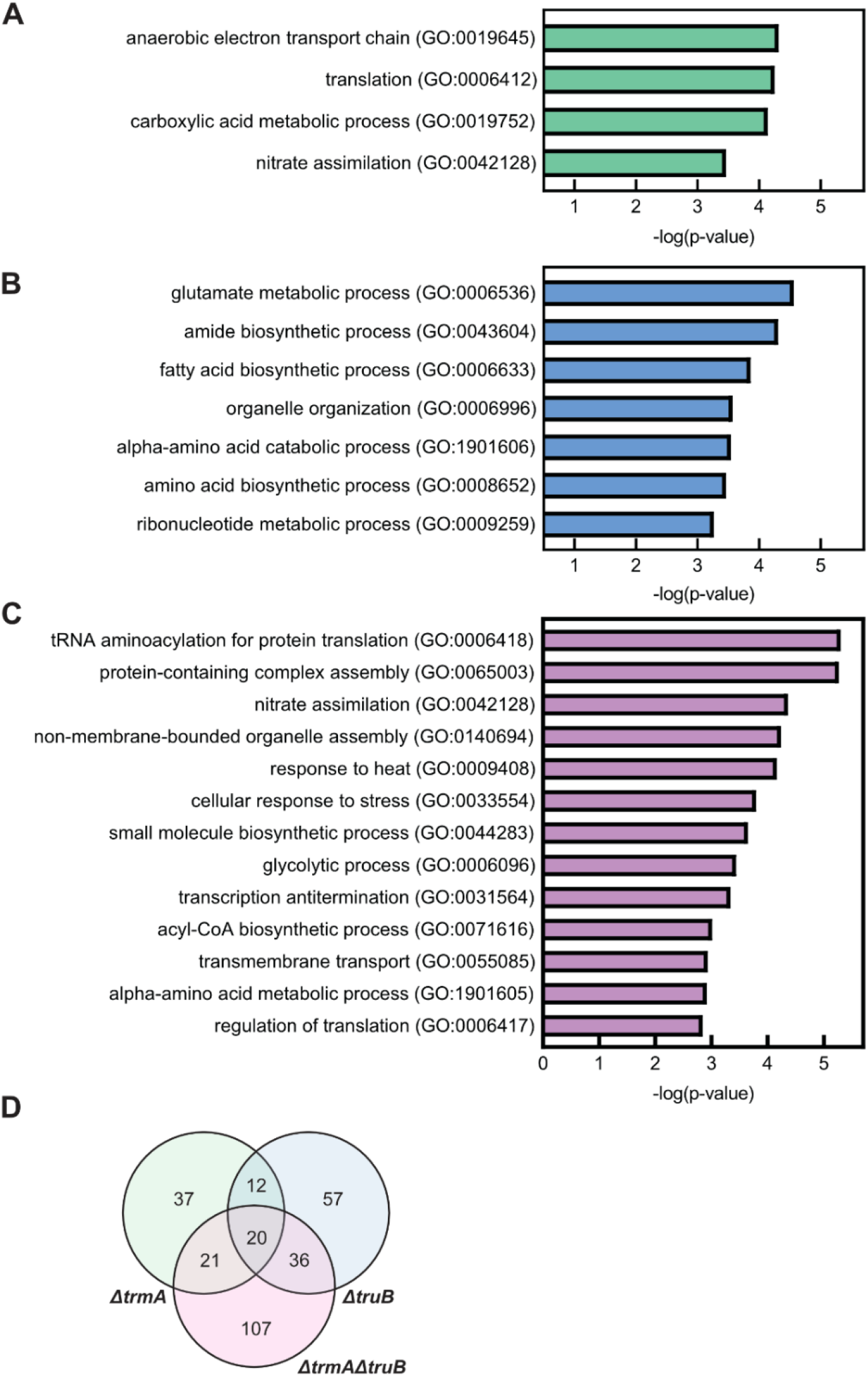
Gene ontology analysis for up/downregulated proteins in *ΔtrmA* (A)*, ΔtruB* (B), and *ΔtrmAΔtruB* (C) strains and overlap between dysregulated proteins for each strain (D).

**Table S1.**
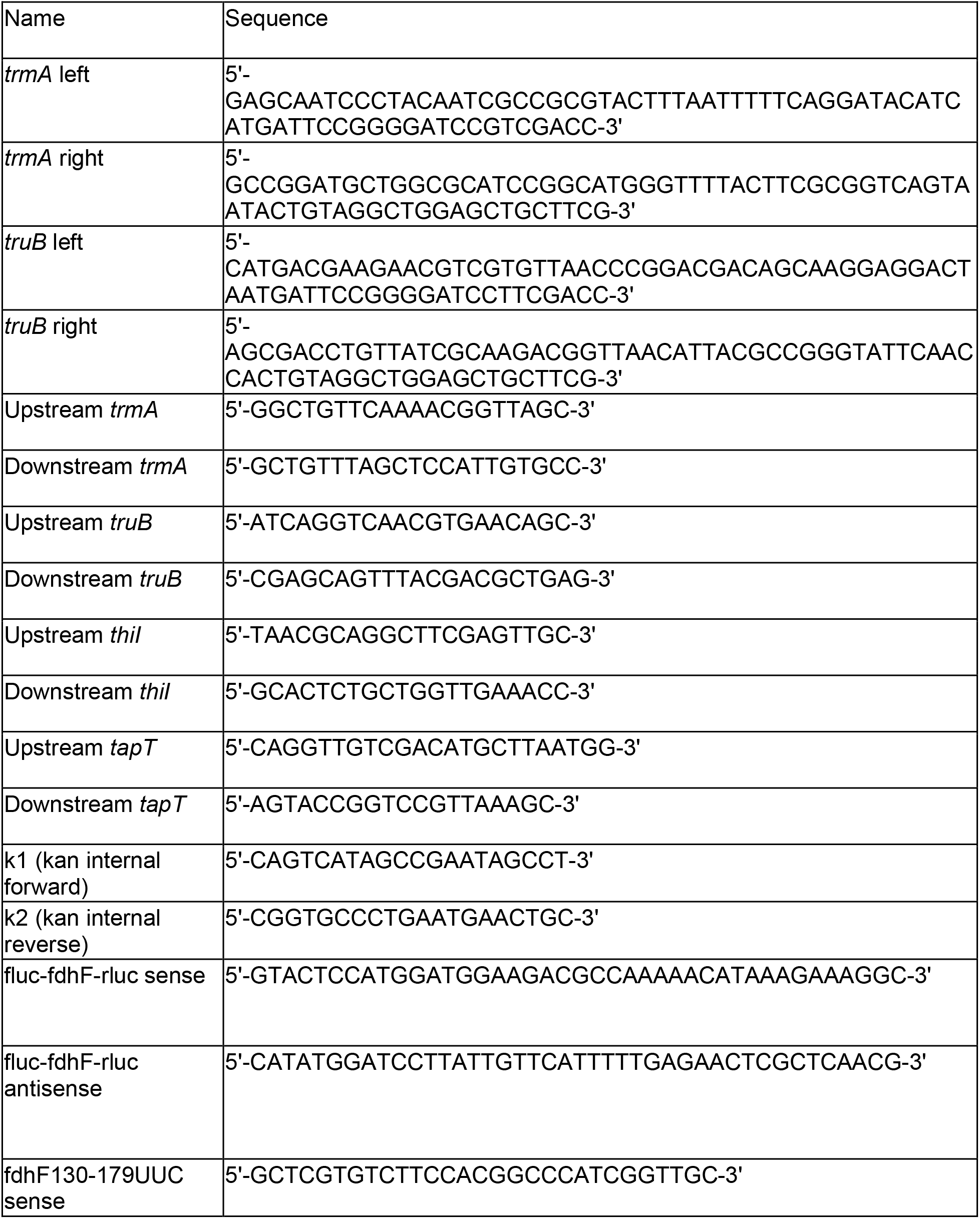
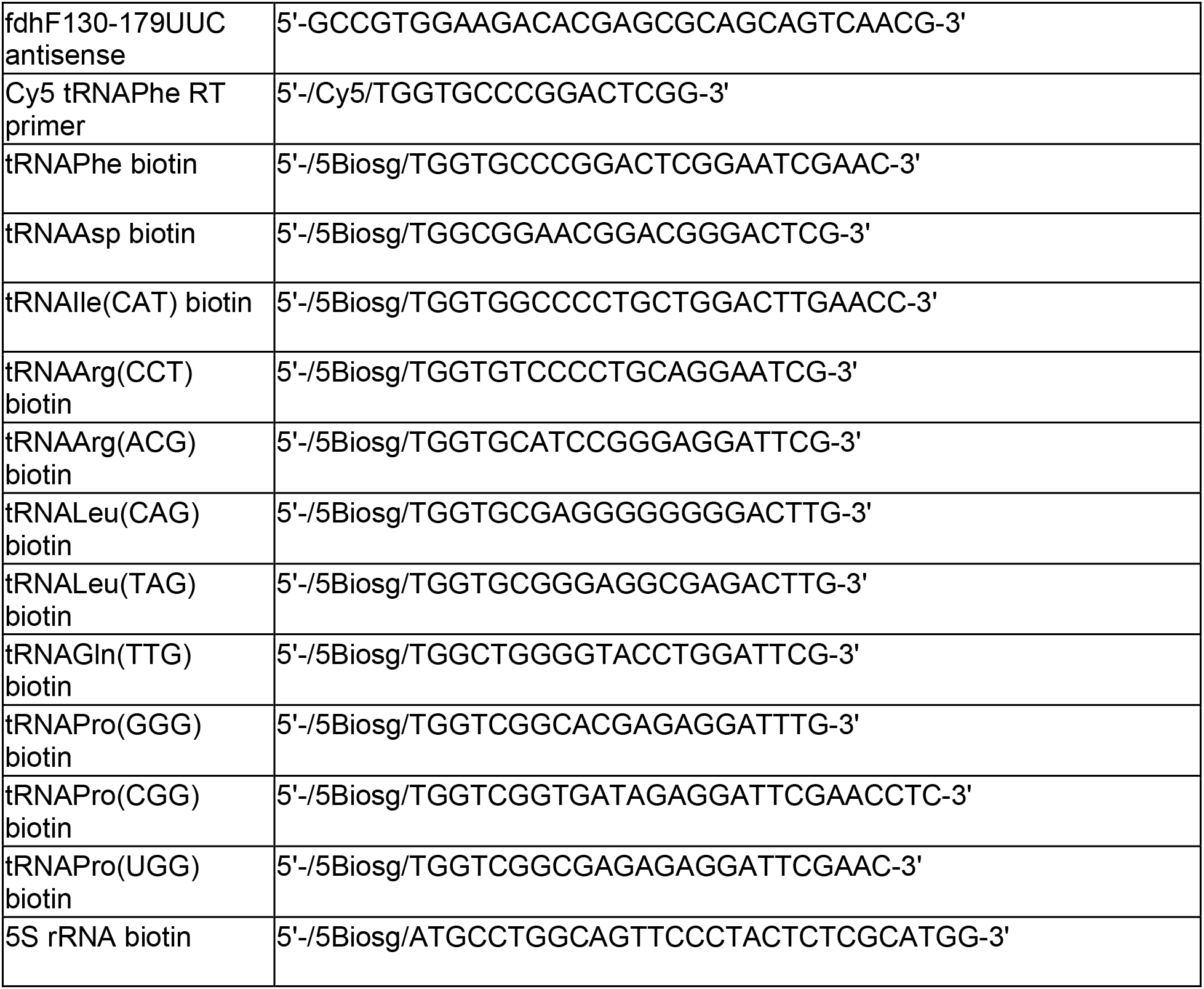
Oligonucleotides used in this study (primers and Northern blotting probes)

## Notes

### Competing Interest Statement

The authors have declared no competing interest.

